# A mechanism for exocyst-mediated tethering via Arf6 and PIP5K1C driven phosphoinositide conversion

**DOI:** 10.1101/2021.10.14.464363

**Authors:** Hannes Maib, David H. Murray

## Abstract

Polarized trafficking is necessary for the development of eukaryotes and is regulated by a conserved molecular machinery. Late steps of cargo delivery are mediated by the exocyst complex, which integrates lipid and protein components to tether vesicles for plasma membrane fusion. However, the molecular mechanisms of this process are poorly defined. Here, we reconstitute functional octameric human exocyst, demonstrating the basis for holocomplex coalescence and biochemically stable subcomplexes. We determine that each subcomplex independently binds to phosphatidylinositol 4,5-bisphosphate (PI(4,5)P_2_), which is minimally sufficient for membrane tethering. Through reconstitution and epithelial cell biology experiments, we show that Arf6-mediated recruitment of the lipid kinase PIP5K1C rapidly converts phosphatidylinositol 4-phosphate (PI(4)P) to PI(4,5)P_2_, driving exocyst recruitment and membrane tethering. These results provide a molecular mechanism of exocyst-mediated tethering and a unique functional requirement for phosphoinositide signaling on late-stage vesicles in the vicinity of the plasma membrane.

## Introduction

Metazoan cell biology is characterized by directional membrane trafficking pathways supporting membranous organelles. These structures are interconnected, with logistics facilitated by both vesicular transport mechanisms and direct membrane contact sites. Strikingly, the specialization of organelles and their distinct identities are maintained, a result of continuous sorting and exchange of phospholipid and protein content. In the polarized trafficking pathway, this sorting includes polarity determinants destined to the apical or basal domains of epithelia, integrins and metalloproteases to the leading edge of invasive cancers, and the transport of receptors to post synaptic densities in neurons (*1*, *2*). The diversity of these cargos highlights that the molecular mechanisms of polarized trafficking and its regulation are fundamental to a broad swath of cell biology, including organismal development (*3*), tissue integrity (*1*), and viral egress (*4*). Importantly, these distinct cargo carriers with diverse cellular functions are all ultimately trafficked to, and fuse with, the plasma membrane in the final step of their journey. However, while the molecular basis for the final step of membrane fusion is well-established, a generalizable mechanistic model for the delivery of these diverse cargo carriers remains outstanding.

Orchestration of polarized trafficking requires a highly conserved trafficking machinery. Early screens for factors responsible for polarized secretion in yeast identified the components of the exocyst complex and its regulatory machinery, including small GTPases (*5*). The connection of this complex to plasma membrane targeting of cargo vesicles in both yeast and mammalian cells is well established (*6*, *7*). Yet, models for exocyst-dependent trafficking in metazoan invoke diverse mechanisms, including GTPase-mediated assembly (*8*), interconnections with the SNARE fusion machinery (*9*, *10*), and membrane-associated cellular factors (*11*–*13*). Exocyst-regulatory factors are linked to phosphoinositide signaling, most notably evidenced by Rab11 (*14*) and Arf6 (*15*–*19*). Indeed, exocyst conservation (*20*) and connection of lipid kinases to polarized secretion (*21*–*25*) suggest a similar interdependence of phosphoinositides and exocyst function.

Phosphoinositides are a family of phospholipids that are signposts for membrane identity (*26*). Their interconversion by lipid kinases and phosphatases is crucial during membrane trafficking and especially well characterized in the endo/lysosomal system (*27*). At the plasma membrane, PI(4,5)P_2_ is enriched and important for cargo delivery and fusion (*15*, *28*, *29*). However, the membrane identity of cargo vesicles directly prior to plasma membrane delivery is poorly defined. Importantly, phosphatidylinositol-4 phosphate (PI(4)P) is established on vesicles originating from both the Golgi and recycling endosomes (*30*–*32*). Furthermore, the delivery of cell polarity proteins to the plasma membrane, such as ß-integrin and e-cadherin, is dependent on the direct recruitment of phosphatidylinositol 4-phosphate 5-kinase type-1 gamma (PIP5K1C), which converts PI(4)P into PI(4,5)P_2_ (*25*, *33*). This dependence implies that the phosphoinositide composition of some cargo vesicles is converted to PI(4,5)P_2_ along the route to the plasma membrane (*32*). Indeed, this outcome has been directly observed during the polarized trafficking of Par proteins during oogenesis in *Drosophila* (*34*). However, no mechanism is known by which PI(4,5)P_2_ positive trafficking vesicles are tethered to the plasma membrane (*35*).

The exocyst is a multi-subunit tethering complex (*2*) formed from eight homologous subunits: Exoc1-8 in mammals and Sec3, Sec5, Sec6, Sec8, Sec10, Sec15, Exo70 and Exo84 in yeast. Importantly, two distinct subunits bind to PI(4,5)P_2_ (*36*, *37*). The binding sites are located at a conserved pleckstrin homology (PH) domain in Exoc1 and in a polybasic region of Exoc7. All exocyst subunits bear a coiled-coil followed by an alpha-helical repeat region (*38*). Through antiparallel coiled-coil assembly, two exocyst heterotetrameric subcomplexes form, with Exoc1-4 as part of Subcomplex-1 and Exoc5-8 in Subcomplex-2 (*38*–*40*). The coiled-coil assemblies of these subcomplexes are analogous to the parallel helical bundle of the soluble n-ethylmaleimide-sensitive-factor attachment receptor (SNARE) membrane fusion machinery (*41*). The stability of coiled-coil structures (*42*) and the *in vivo* colocalization of exocyst subunits (*43*, *44*), suggests that the exocyst is present as either tetrameric subcomplexes, or a full octamer. Indeed, each subcomplex harbors a distinct PI(4,5)P_2_ binding site, though the function of this duplication is unknown.

The interconnectivity of exocyst subunits enables a potential for regulatory modification through disassembly of the holocomplex. Indeed, live cell imaging of endogenously tagged exocyst subunits suggests that although the subcomplexes are recruited to vesicles independently, they arrive at the PM together (*43*). Furthermore, deletion of one single subunit leads to a concomitant decrease in vesicle tethering (*36*, *43*), highlighting the full holocomplex as the functional unit for vesicle tethering with the PM. However, a common mechanism for exocyst-mediated tethering of cargo carriers to the plasma membrane is elusive.

Here, we reconstitute the complete human exocyst complex from its individual subunits to address the mechanism by which it tethers vesicles to the plasma membrane during polarized trafficking. We find that the complete octamer readily associates from its separate subunits and is suborganized into two distinct subcomplexes with differing modes of assembly. Further, we show that each of these subcomplexes independently binds to PI(4,5)P_2_ and that the holocomplex tethers two opposing PI(4,5)P_2_-positive membranes. Using cryo-electron tomography, we directly visualize this exocyst-dependent tethering of liposomes and determine a regular tethering distance consistent with the known dimensions of the exocyst complex. Next, we determined how exocyst function is linked to PI(4,5)P_2_. *In vitro* reconstitution of phosphoinositide conversion by PIP5K1C is sufficient to drive both exocyst recruitment and membrane tethering. Using endogenous knock-in cell lines, we observe exocyst colocalization with PIP5K1C at dynamic subdomains of the plasma membrane. Finally, we demonstrate that PI(4)P to PI(4,5)P_2_ conversion is driven by Arf6 activation both *in vitro* and in cells. Therefore, we propose a model in which phosphoinositide conversion, mediated by Arf6 and PIP5K1C, regulates exocyst-dependent membrane tethering.

## Results

### Complete human exocyst connectivity and reconstitution

To understand the mechanism of exocyst holocomplex assembly and tethering, we reconstituted the human exocyst complex using baculovirus-mediated insect cell expression (Fig. 1) (*45*). We co-expressed every combination of exocyst subunits in pairs, with one subunit harboring a single GST-tag for isolation (Fig. 1A). Subsequently we performed pulldowns of the GST-tagged bait subunit to determine if the prey is directly isolated in this overexpression approach. We used SDS-PAGE followed by Coomassie staining to identify binary interaction partners between the subunits of the exocyst holocomplex (Fig. 1B), resulting in several apparent dimers. Next, using the information garnered by the dimer analysis, we co-expressed and analyzed the logical trimers (Fig. 1C), determining that Exoc7 binding requires the formation of an initial Exoc5:6 dimer. Subsequently, because we observed no interaction of Exoc8 in the trimeric interaction experiment (Fig. 1C), we isolated a trimer of Exoc5:6:7 and confirmed its binding to Exoc8 (Fig. 1D). Most importantly, deriving from these results are two subcomplexes, named Subcomplex-1 (Exoc1:2:3:4) and Subcomplex-2 (Exoc5:6:7:8). These differ in their assembly, with Subcomplex-1 based upon binary subunit interactions, and for Subcomplex-2, a hierarchal assembly. Building on the initial formation of an Exoc5:6 dimer, Exoc7 and Exoc8 are subsequently recruited to form tetrameric Subcomplex-2.

**Fig. 1.**
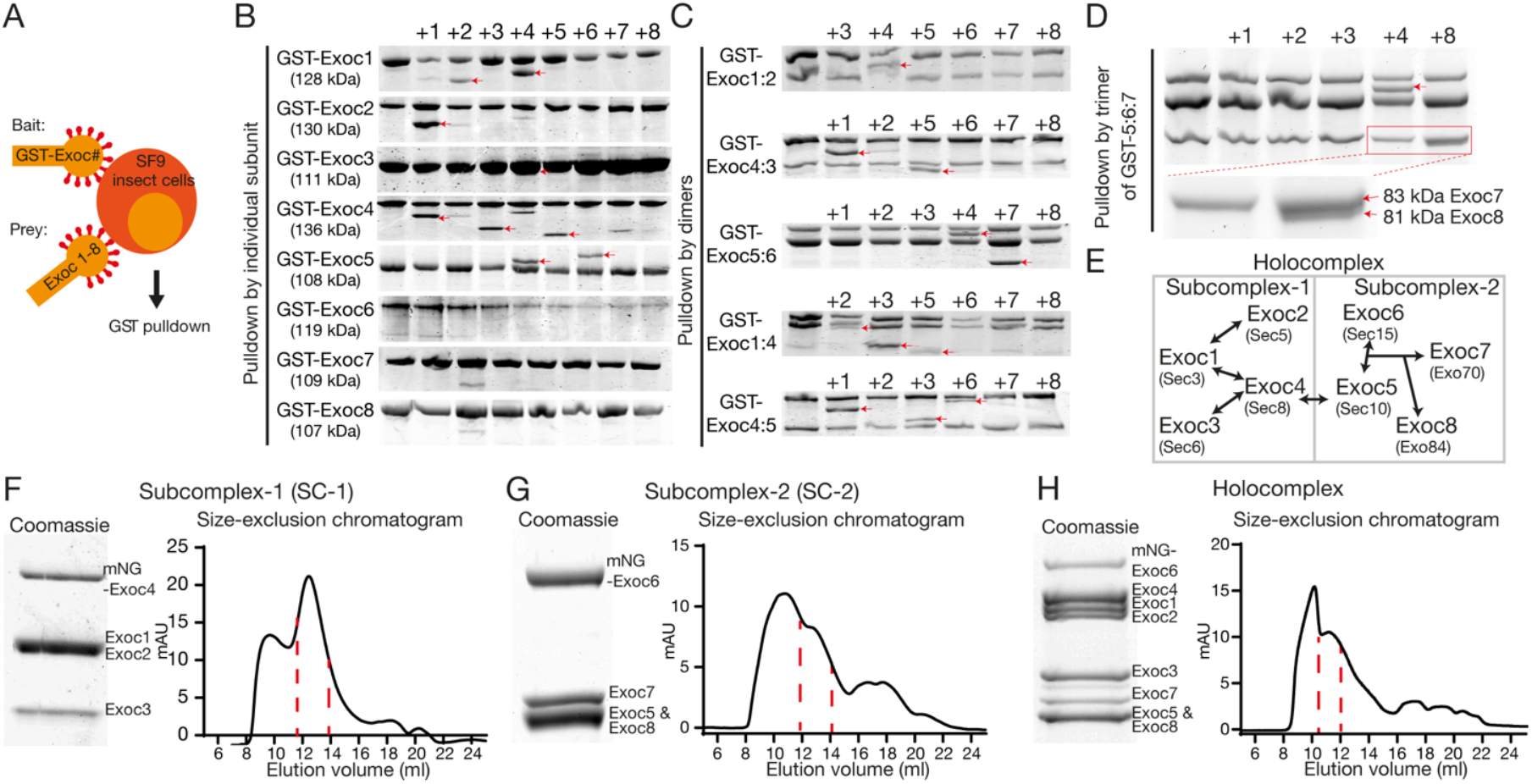
Subunit connectivity and reconstitution of complete human exocyst complex. **A**, Schematic of subunit interaction screen. SF9 insect cells were co-infected with baculovirus encoding single GST-tagged subunits of the exocyst complex (bait) together with untagged subunits (prey), and analyzed via GST pulldowns followed by SDS-PAGE and Coomassie staining. **B,C**, Dimeric and trimeric interactions. First lanes are GST-bait without prey. Numbers above gel bands indicate each of the coexpressed exocyst subunits 1-8. Interaction partners are highlighted with red arrowheads. **D**, Tetrameric interactions. GST-Exoc5 together with Exoc6 and Exoc7 were coexpressed with the remaining subunits. **e)** Schematic of complete human exocyst subunit connectivity. Holocomplex is divided into Subcomplex-1 and Subcomplex-2, which are joined by an interaction of Exoc4 with Exoc5. Names of yeast subunits are given for reference. **F-H**, Biochemical reconstitution of exocyst Subcomplex-1, −2 and holocomplex with size-exclusion chromatograms (Superose 6 Increase 10/300 GL; void volume, 9 mL). Both Subcomplex-1 and 2 elute at a retention volume of 13 ml while full holocomplex elutes at 11 ml. Coomassie-stained insets are taken from peak fractions. Full gels are found in Fig. S4-5.

To assess the assembly of pentamers to octamers, we separately expressed individual subunits and mixed them upon lysis (Fig. S1A-C). The results from the subsequent pulldowns enabled us to conclude that exocyst assembly does not require co-translation nor input *in vitro* from small GTPases. Moreover, the human exocyst efficiently self-assembles in a hierarchal manner, evolutionarily conserved from yeast (Fig. 1E) (*38*–*40*, *46*). The connection between Subcomplex-1 and −2 is dependent on a strong interaction between Exoc4 and Exoc5, resulting in holocomplex assembly.

The principles of assembly for exocyst intermediates and subcomplexes enabled us to discern a purification strategy for intact complexes. For functional analysis, we included fluorescent tags in positions structurally and empirically determined appropriate (Fig. 1F-H) (*47*). An assembly mechanism driven by the largely bipartite CorEx regions, and stabilized by other multivalent interactions (*46*), enabled us to identify the subunits necessary for purification of the holocomplex (*48*). Purified holocomplex is stoichiometric, as determined by quantitative mass spectrometry and supported by native PAGE (Fig. S1H,I). Moreover, negative stain electron microscopy revealed individual isolated complexes, and processing by single-particle methods results in a low-resolution model with dimensions consistent with that derived from cryo-electron microscopy of the yeast complex (Fig. S1J). Taken together with the rules for its assembly, reconstitution of the human exocyst opens a unique opportunity to gain mechanistic insight into its function.

### Two independent PI(4,5)P_2_ binding sites enable exocyst membrane tethering

Key to exocyst function is its ability to bind lipids. A late step in delivery of some cargoes to the plasma membrane is the conversion of a subset of vesicular phosphoinositides to PI(4)P (*30*, *31*). This change of lipid chemistry is concomitant with trafficking of vesicles destined for the plasma membrane that harbor cargos critical for polarity maintenance and cellular homeostasis (*34*, *35*, *49*, *50*). Exoc1 and Exoc7 bind phosphoinositides in isolation (*36*, *37*), but cooperativity or independence in membrane binding has not been assessed.

To determine the selectivity of exocyst complexes to phosphoinositides, we measured the binding of purified, fluorescently-tagged holo- and subcomplexes for PI(4)P or PI(4,5)P_2_ in membranes chosen to mimic the plasma membrane. We formed supported-lipid bilayers on silica beads with liposomes composed of 85% 1-palmitoyl-2-oleoyl-glycero-3-phosphocholine (POPC), 10% phosphatidylserine and 5% phosphoinositide (Fig. 2A) (*51*, *52*). By including N-terminal mNeonGreen (mNG) in Exoc4 for Subcomplex-1, and Exoc6 for both Subcomplex-2 and the holocomplex, we observed recruitment of the complexes to membrane-coated beads by confocal microscopy (Fig. 2B). To statistically compare exocyst recruitment, we segmented and quantified the integral fluorescence intensity of the bound exocyst per membrane-coated bead (Fig. 2C) (POPC/PI(4)P versus PI(4,5)P_2_; p<0.01). The exocyst and Exoc1/Exoc7-containing subcomplexes are unambiguously recruited selectively to PI(4,5)P_2_ and not PI(4)P-containing membranes at non-saturating conditions (100 nM complex). Moreover, by harnessing the assembly hierarchy to generate exocyst subcomplexes, we explicitly demonstrate the requirement for Exoc1 or Exoc7 subunits for membrane binding (Fig. S2) (*35*). Strikingly, the novel observation that both Subcomplex-1 and −2 independently bind PI(4,5)P_2_, and not PI(4)P, questions the potential role of avidity in membrane binding by the holocomplex.

**Fig 2.**
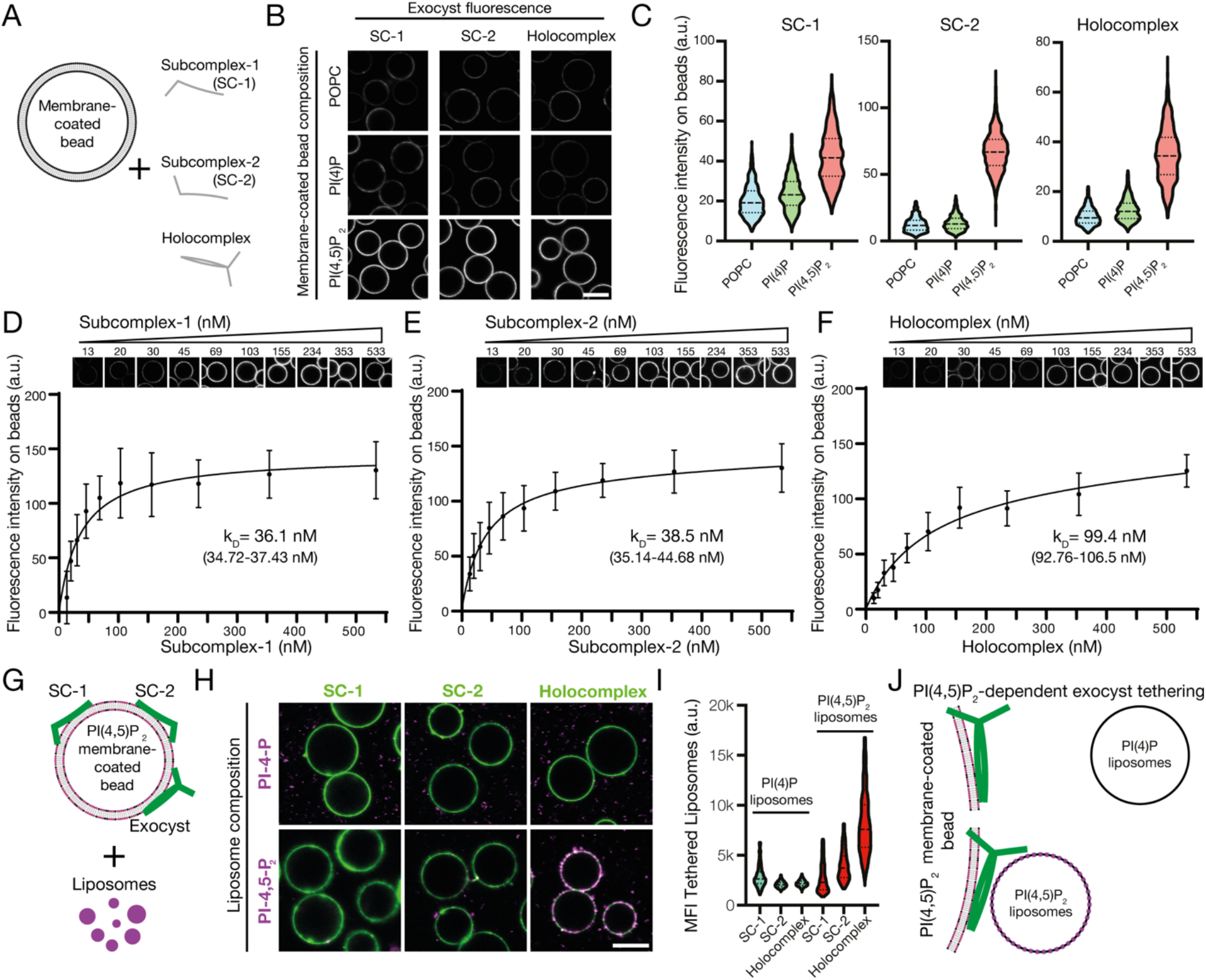
Independent binding of exocyst Subcomplex-1 and −2 to PI(4,5)P_2_ enables holocomplex tethering. **A**, Schematic of phosphoinositide specificity experiment. Membrane-coated beads of distinct lipid composition were generated. To these beads, fluorescently-tagged exocyst subcomplexes were added to assess membrane binding by fluorescence microscopy. **B**, Reconstituted fluorescently-tagged Subcomplex-1, −2, or holocomplex were added at 100 nM to membrane-coated beads. These were formed from a lipid composition of 84.9% 1-palmitoyl-2-oleoyl-glycero-3-phosphocholine (POPC), 10% phosphatidylserine and 5% phosphoinositide or POPC, doped with 0.1% rhodamine-DPPE. Beads were imaged by confocal microscopy. **C**, Membrane-coated beads were segmented using the rhodamine lipid signal as a membrane mask, and the mean fluorescence intensities of exocyst complexes bound to each bead were determined. Violin plots, 877-1395 segmented beads per condition. **D-F**, Relative PI(4,5)P_2_ membrane binding affinities for the exocyst complexes. Subcomplex-1, −2, or holocomplex were added in a dilution series to membrane-coated beads containing 5% PI(4,5)P_2_ and imaged by confocal microscopy. Individual beads were segmented, and a binding saturation curve was fitted resulting in mean kD determination with 95% confidence intervals given. 340-1359 beads were quantified per concentration, mean±standard deviation. **G**, Schematic of liposome tethering experiment. Subcomplex-1, −2, or holocomplex (150 nM) were bound to 5% PI(4,5)P_2_ membranes. Liposomes doped with Atto647N-DOPE and composed with either 5% PI(4)P or PI(4,5)P_2_ were added. **H-I**, Membranes and exocyst complexes were imaged by confocal microscopy, segmented and the fluorescent intensity of tethered liposomes was measured. Violin plots from 430-778 individual beads. MFI, mean fluorescent intensity. **J**, Exocyst holocomplex tethers two PI(4,5)P_2_, but not PI(4)P, containing membranes to each other.

To quantitate the independence and modality of lipid binding, we determined the relative affinities of Subcomplex-1, −2, and the holocomplex for PI(4,5)P_2_ membranes. We adapted the membrane-coated bead assay with varying protein concentrations to obtain their relative affinities for PI(4,5)P_2_. By systematic imaging and analysis, we determined that both Subcomplex-1 and −2 were recruited to membranes with similar strength (Subcomplex-1, *K_D,relative_*=36.1 nM; Subcomplex-2, *K_D,relative_*=38.5 nM) (Fig. 2D,E). However, the exocyst holocomplex bound to membranes of identical composition significantly weaker (*K_D,relative_*=99.4 nM; AIC<0.01%) (Fig. 2F). Interestingly, this result makes monovalent binding to PI(4,5)P_2_ by the exocyst plausible, though with an increased kinetic off rate compared to individual subcomplexes. Moreover, it negates the possibility of avidity in lipid-mediated recruitment of the holocomplex to membranes. These results, together with phosphoinositide selectivity (Fig. 2B, C; Fig. S2) and the holocomplex assembly mechanics (Fig. 1), suggest exocyst lipid binding occurs in a *trans* configuration. Therefore, we hypothesized that the holocomplex is capable of tethering two distinct PI(4,5)P_2_-containing membranes.

To test this hypothesis, we recruited the fluorescently-tagged exocyst or subcomplexes (150 nM) to PI(4,5)P_2_-containing POPC membrane-coated beads (Fig. 2G) as previously demonstrated (Fig. 2B). Next, we added fluorescently-labeled liposomes harboring either PI(4)P or PI(4,5)P_2_ and quantified the fluorescence intensity of tethered liposomes (*51*). Membranes were only tethered in the presence of the exocyst holocomplex when both contained PI(4,5)P_2_ (Fig. 2H). Individual exocyst subcomplexes failed to tether liposomes, and in the absence of liposomal PI(4,5)P_2_, we observed no statistically significant tethering (Fig. 2I). These data provide unequivocal evidence for a minimal mechanism of exocyst-mediated membrane tethering (Fig. 2J).

To verify these findings, and to directly observe the exocyst holocomplex on membranes, we harnessed cryo-electron tomography. We prepared samples of 5% PI(4,5)P_2_ - containing POPC liposomes extruded to 100 nm and mixed with exocyst. Samples were plunge-frozen, micrographs acquired, and tomograms reconstructed to visualize the samples in three dimensions. In tomograms, we observed a regular proteinaceous material, disperse in regions devoid of liposomes. In regions with liposomes, we observed membrane recruitment of the exocyst complex, and tethering of membranes (Fig. 3). By manual segmentation of the tomograms in three-dimensions, we noted and quantified a regular distance between pairs of membranes at the position of closest approach. This resulted in an average distance of 32.1 nm, consistent with the dimensions of the exocyst complex from yeast (*38*), and providing additional evidence for exocyst-mediated tethering of PI(4,5)P_2_ membranes.

**Fig.3.**
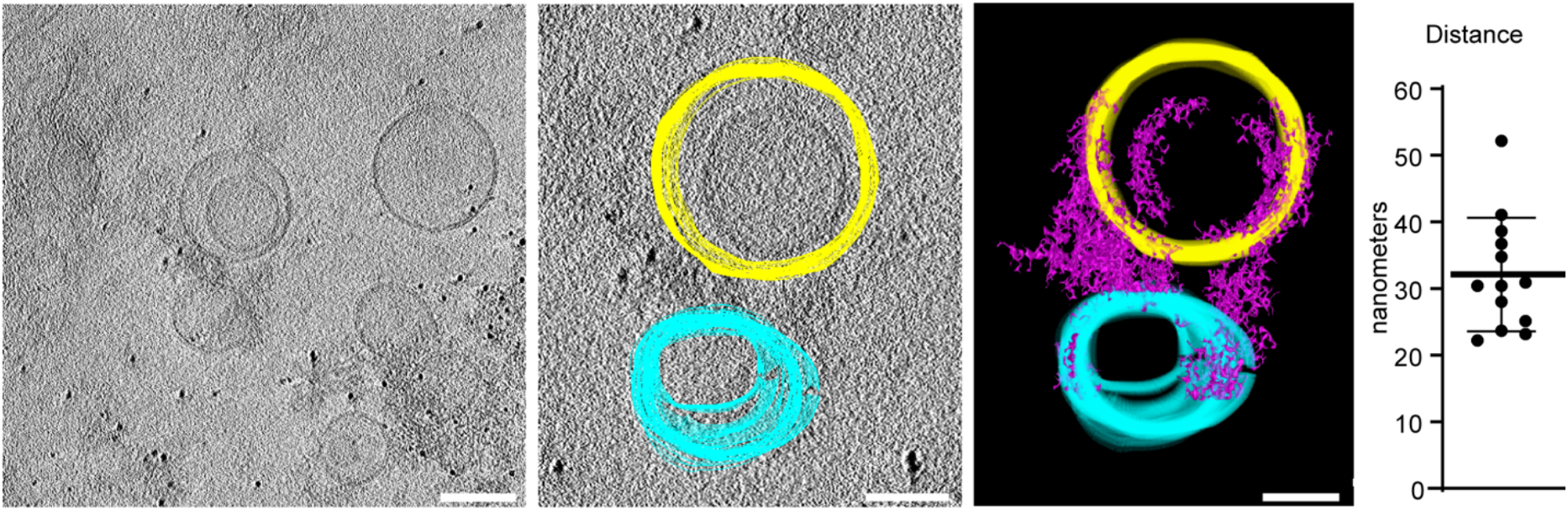
Cryo-electron tomography of exocyst-mediated liposome tethering. Purified exocyst holocomplex was added to 5% PI(4,5)P_2_ containing liposomes in the presence of 5 nm Protein-A coated gold fiducial markers, plunge frozen in liquid ethane and tilt series were acquired. Tomograms were reconstructed resulting in a three-dimensional data set, and liposomes segmented manually through the z-axis slices (yellow and cyan). Exocyst holocomplex, present as proteinaceous material bound to the liposomes, was segmented using auto-thresholding (purple), which also partially delineates electron-dense portions of the liposomes. A model was generated from this segmentation, and the distance of closest approach between tethered liposome pairs was determined (mean ± standard deviation). Scale bars, 100 nm.

### Phosphoinositide conversion drives membrane tethering

In polarized trafficking, phosphoinositide dynamics are required for delivery of vesicles prior to the engagement of fusion machinery. A key factor in these dynamics is the endosomal recruitment of the phosphatidylinositol-5 kinase PIP5K1C (*18*, *30*, *33*). To determine if phosphoinositide conversion drives exocyst-mediated tethering, we reconstituted human PIP5K1C to convert PI(4)P into PI(4,5)P_2_ in a modified assay for exocyst recruitment (Fig. 4A) (*53*). We mixed recombinant fluorescently-tagged exocyst (100 nM) with PI(4)P-positive membrane-coated beads (as in Fig. 2B). After the addition of reconstituted PIP5K1C (150nM), we observed exocyst recruitment to membranes in an ATP-dependent manner by confocal fluorescence microscopy (Fig. 4B). In this system, limited by mixing, the conversion and subsequent exocyst recruitment occurred rapidly once underway. This recruitment is consistent with the observed affinities of the exocyst to PI(4,5)P_2_ membranes (Fig 2D-F) and provides a baseline for the activity of PIP5K1C, in the absence of a mechanism for lipid kinase recruitment. Critically, phosphoinositide conversion not only drove exocyst membrane recruitment, but also tethering. We mixed PI(4,5)P_2_-positive liposomes, the exocyst, and PIP5K1C together with PI(4)P-positive membrane-coated beads. Following conversion until a saturating time point, we evaluated the fluorescence intensity of tethered liposomes (Fig. 4C,D). We observed a clear dependence of liposome tethering on phosphoinositide conversion, and a linear dependence on the amount of exocyst recruited. Therefore, *de novo* generation of PI(4,5)P_2_ by PIP5K1C mediates *in vitro* exocyst-dependent tethering. This suggests that late-stage phosphoinositide conversion drives cargo vesicle tethering during polarized trafficking.

**Fig. 4.**
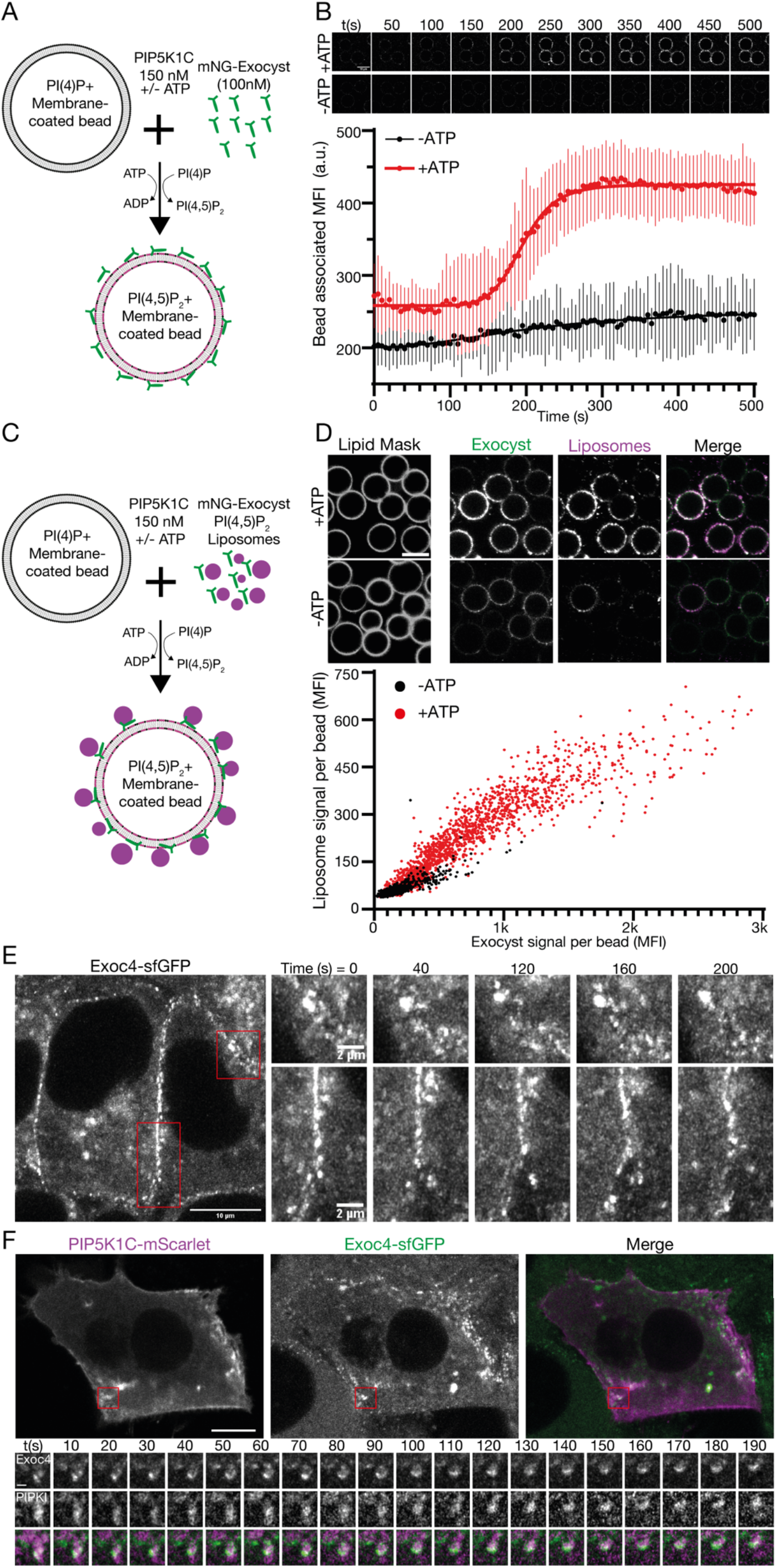
Phosphoinositide conversion drives exocyst-mediated membrane tethering. **A**, Schematic of exocyst recruitment by phosphoinositide conversion. Membrane-coated beads containing PI(4)P were mixed with exocyst (100 nM) and PIP5K1C (150 nM) in the presence and absence of ATP. **B**, Exocyst is recruited to membrane-coated beads upon ATP-dependent PI(4)P to PI(4,5)P_2_ conversion, and imaged by confocal microscopy. Beads were segmented, and the associated exocyst mean fluorescent intensity was measured over time. Mean ± standard deviation, n=44, 68 beads. Scale bar 10 μm. **C**, Schematic of membrane tethering by phosphoinositide conversion. PI(4,5)P_2_ containing liposomes were added to the phosphoinositide conversion experiment. **D**, The intensity of bound exocyst and tethered liposomes was determined after 30 minutes, and the mean fluorescent intensity of each were plotted per bead. n=1440, 1864 beads. Scale bar, 10 μm. **E**, NMuMG cells with endogenous Exoc4-sfGFP were imaged by livecell Airyscan microscopy. Insets show the dynamics of plasma membrane-associated exocyst subdomains. **F**, Cells were transfected with mScarlet-tagged PIP5K1C, and imaged by live-cell Airyscan microscopy. Inset shows time series of exocyst colocalization with PIP5K1C at dynamic subdomains of the plasma membrane. Scale bar, 10 μm, inset scale bar, 1 μm.

To validate these findings in a cellular system, we used genome edited knock-in cells from mouse mammary epithelia (NMuMG) with sfGFP at the C-terminus of Exoc4 (*43*). These cell lines have previously been described, and the fluorescent tag does not affect exocyst function. Importantly, these cells readily differentiate into an epithelial sheet with apicobasal polarity and are therefore well-suited to observe the exocyst during polarized trafficking. Using live-cell Airyscan microscopy (*54*), we determined that the exocyst is mainly located at basolateral and junctional membranes. Here, the exocyst is organized into dynamic subdomains that are stable for several minutes (Fig. 4E). This localization is consistent with its role as a tethering complex in the direct vicinity of the plasma membrane (*2*, *43*). To address the link between phosphoinositide conversion and exocyst function, we exogenously expressed PIP5K1C-mScarlet-I in these cell lines and performed live cell imaging (Fig 4F). We observed PIP5K1C and the Exoc4-sfGFP together at dynamic subdomains in the direct vicinity of the plasma membrane. This data provides further evidence for a functional link between the exocyst and the generation of PI(4,5)P_2_ (*33*).

### Arf6 mediates PIP5K1C-dependent PI(4,5)P_2_ generation and exocyst recruitment

The conversion of PI(4)P into PI(4,5)P_2_ at intracellular membranes is catalyzed by the Arf6-mediated recruitment of PIP5K1C (*25*), which is in part also dependent on the presence of anionic lipids (*53*). We hypothesized that regulation of Arf6, by virtue of recruiting PIP5K1C, drives PI(4,5)P_2_ generation, and subsequent exocyst-mediated tethering. To observe cellular phosphoinositides, we harnessed a staining approach using purified phosphoinositide-binding probes, allowing us to visualize lipid populations in fixed cells (Fig. S3A, B) (*55*). Briefly, recombinant purified SidC (*56*), and PLCδ-PH domain (*57*), tagged with GFP were used as probes in fixed and permeabilized NMuMG cell lines to visualize PI(4)P and PI(4,5)P_2_, respectively. We observed strong colocalization of immunostained Golgi-resident GM130 with PI(4)P by SidC staining (Fig. S3A,B). Respectively for PLCδ-PH staining, we observed plasma membrane alongside intracellular vesicles harboring PI(4,5)P_2_ and e-cadherin in PIP5K1C expression conditions (Fig S3B). This methodology allows us to robustly detect phosphoinositides in cells while avoiding overexpression artifacts of lipid probes.

To address Arf6-mediated regulation of the conversion of PI(4)P to PI(4,5)P_2_, we used expression of constitutively active or inactive Arf6 to disrupt GTPase cycling (Arf6-Q67L and Arf6-T27N, respectively) (*58*). Expression of Arf6-T27N results in the well-described phenotypes of post-Golgi halted secretion, while expression of Arf6-Q67L lead to a progressive accumulation of plasma membrane-derived vesicles resulting from overactive macropinocytosis (e.g. Fig. 5A, top left panel) (*15*, *59*, *60*). We took advantage of these phenotypes to determine the localization of phosphoinositides to the resulting internal membranes. Arf6-T27N positive membranes contained PI(4)P (Fig. 5A) but not PI(4,5)P_2_ (Fig. 5B). Moreover, they were only partially positive for a Golgi marker (Fig 3A; top insets), indicating an intracellular PI(4)P pool not associated with the Golgi. In contrast, membranes positive for Arf6-Q67L were strongly enriched with PI(4,5)P_2_ (Fig. 5B), but not PI(4)P (Fig. 5A). The observed differences in colocalization of phosphoinositides with Arf6 correlates directly with its activation state, and is consistent with a direct role in the generation of PI(4,5)P_2_.

**Fig. 5.**
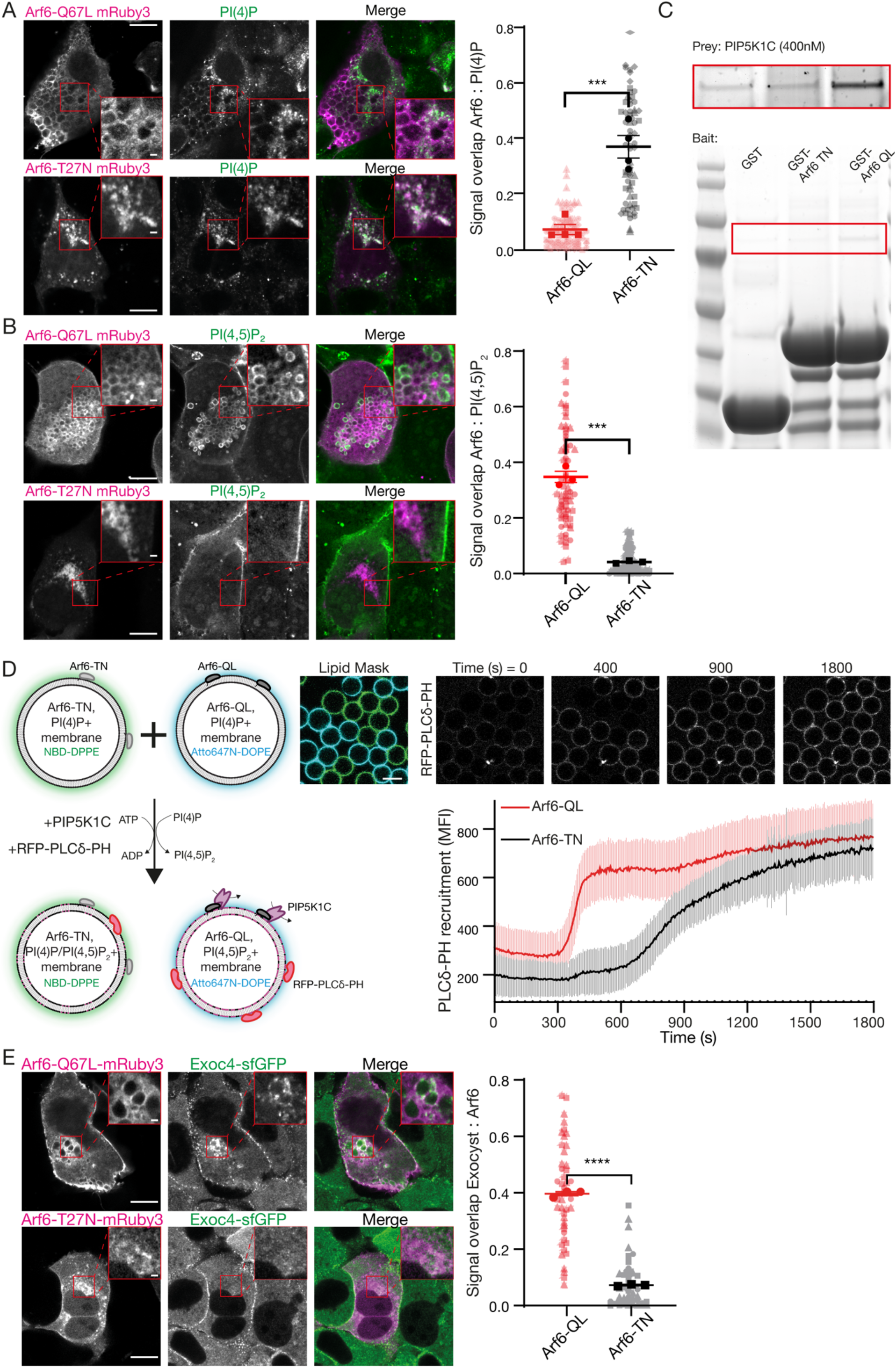
Arf6 controls PI-4,5-P_2_ conversion and exocyst recruitment. **A-B**, NMuMG cells were transfected with Arf6-Q67L or Arf6-T27N tagged with mRuby3, and subsequently fixed, permeabilized and stained against either PI(4)P or PI(4,5)P_2_ using purified SidC or PLCδ-PH fused to GFP, respectively. Signal overlap between channels was determined using the ImageJ plugin Squassh. Plot shows biological repeats (in bold) with individual cells (in background), n=3-4 repeats with 2936 individual cells each. Asterisks denote p<0.001. Scale bar, 10 μm, inset, 1 μm. **C**, Purified GST, GST-Arf6-T27N or GST-Arf6-Q67L were bound to resin as bait, and purified PIP5K1C (400 nM) used as prey. Inset (top) is increased contrast. **D**, Membrane-coated beads harboring PI(4)P and either recombinant Arf6-T27N or Arf6-Q67L were distinctly labelled with NBD-DPPE or Atto647N-DOPE and mixed. PIP5K1C (12.5 nM) was added, and the recruitment of purified PLCδ-PH fused to RFP was measured over time. Beads were segmented using either the NBD or Atto647N signal as masks. Plot is mean fluorescent intensity of recruited PLCδ-PH ± standard deviation, quantified in ImageJ. n=92-99 beads. Scale bars, 10 μm. **E**, NMuMG cells expressing endogenously sfGFP-tagged Exoc4 were transfected with Arf6-Q67L or Arf6-T27N fused to mRuby3 and imaged using Airyscan confocal microscopy. Signal overlap between channels was determined using the ImageJ plugin Squassh. n=3 repeats with 7-25 individual cells each, asterisks denote p<0.001.

To evaluate the mechanism by which Arf6 activation leads to the accumulation of PI(4,5)P_2_, we directly assessed PIP5K1C binding to Arf6 (*16*, *61*). Purified PIP5K1C interacts with purified Arf6 in a GTPase activity-dependent manner (Fig. 5C), consistent with results from cell biological experiments (*15*, *61*) and demonstrating that PIP5K1C is an effector of Arf6 (*16*). This result explains the elevated levels of PI(4,5)P_2_ on intracellular membranes in rf6-Q67L expression (Fig. 5B). Moreover, the direct interaction led us to hypothesize that Arf6-mediated lipid kinase recruitment drives rapid phosphoinositide conversion.

To directly evaluate the biochemical characteristics of Arf6-mediated phosphoinositide conversion in a reduced system, we reconstituted the lipid kinase and membrane-coated beads harboring PI(4)P together with Arf6. We conjugated either Arf6-T27N or Arf6-Q67L to membrane-coated beads harboring the high-affinity TrisNTA-DODA lipid via N-terminal 6xHis tags (*62*). Both populations were distinctly doped with fluorescent lipids and observed simultaneously by microscopy (Fig 5D). After addition of PIP5K1C at kinetic concentrations (12.5 nM) to these membranes, we observed the production of PI(4,5)P_2_, assessed by the recruitment of RFP-PLCδ-PH. As expected, we observed relatively slow recruitment of this probe to Arf6-T27N membranes, reflecting basal activity of the lipid kinase (Fig 5E). In contrast, membranes harboring Arf6-Q67L displayed extremely rapid and cooperative phosphoinositide conversion. This result supports a mechanism whereby active Arf6 potentiates lipid kinase activity by the direct recruitment of PIP5K1C to the membrane. The phosphoinositide conversion thereby functions as a biochemical switch (*63*, *64*), rapidly redefining the identity of the membrane. We demonstrated that this change of identity is sufficient for exocyst recruitment and tethering (Fig. 4), which suggests a key role for phosphoinositide conversion in polarized trafficking.

To directly test this mechanism in a cellular system, we determined the exocyst localization in polarized NMuMG cells expressing Arf6 variants (*43*). In cells expressing the inactive Arf6-T27N, we observed no change to exocyst localization (Fig 5E; compare Fig. 4E). In contrast, and in addition to the normal exocyst localization, expression of active Arf6-Q67L resulted in significant colocalization with the exocyst (Fig 5E), retargeting the complex to PI(4,5)P_2_ positive internal membranes. This result demonstrates that the exocyst is linked to Arf6 in an activity-dependent manner. Therefore, these experiments support a model in which late-stage phosphoinositide conversion is critical to exocyst-mediated tethering in polarized trafficking (Fig. 6).

**Fig. 6.**
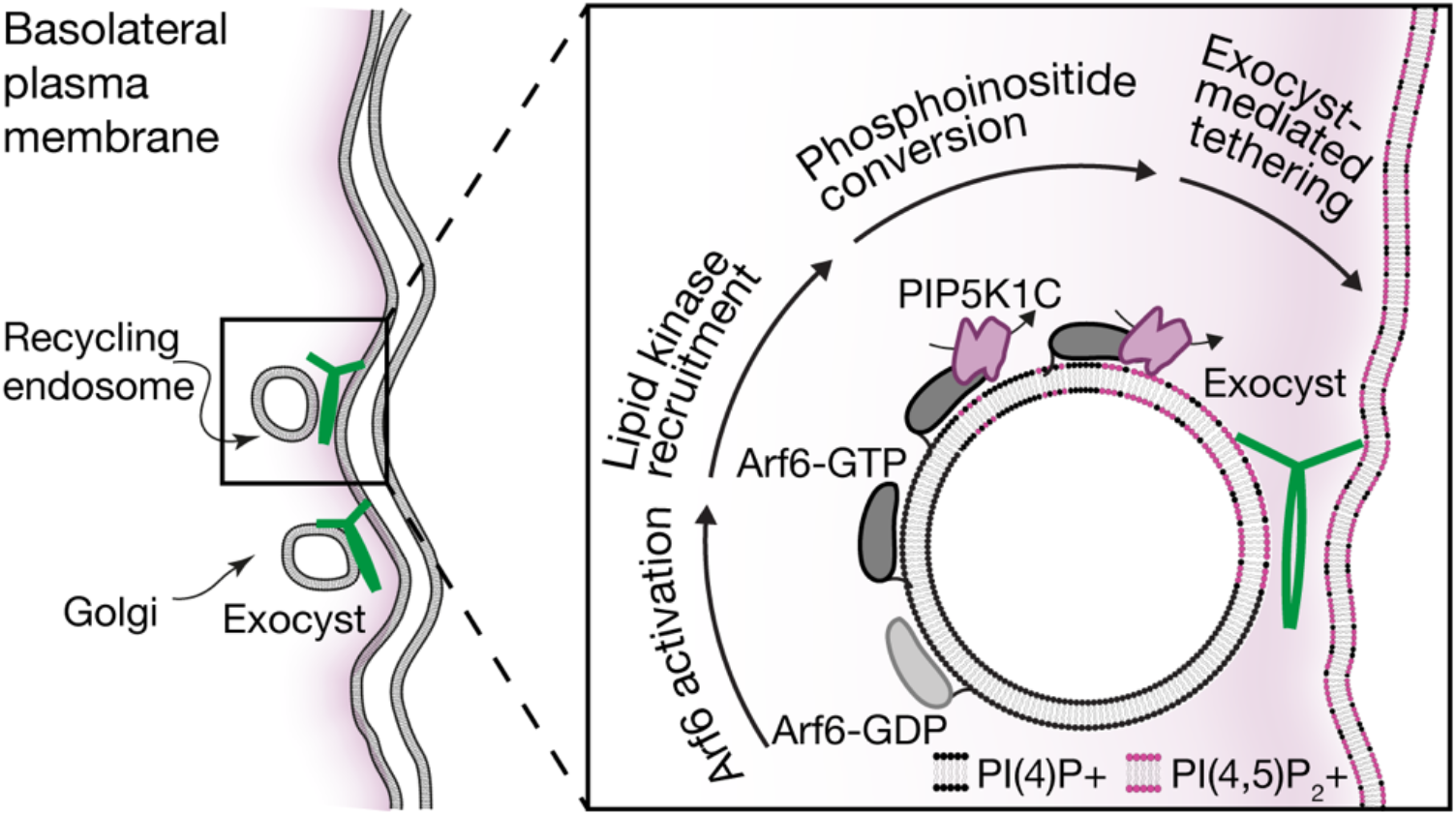
Exocyst-mediated lipid kinase driven membrane tethering. Polarized trafficking vesicles originating either from recycling endosomes or the Golgi are trafficked to the vicinity of the basolateral PM. Close to the PM, Arf6 is activated, resulting in recruitment of the PIP5K1C lipid kinase. This kinase converts vesicular PI(4)P into PI(4,5)P_2_, which results in exocyst recruitment and cargo carrier tethering to the PM.

## Discussion

Exocyst-mediated selectivity and tethering results from the coordination of intracellular PI(4,5)P_2_ conversion at late stages of vesicle delivery to the plasma membrane. We have demonstrated in a reconstitution-based approach the constitutive nature of the exocyst complex, and provide new mechanistic evidence for holocomplex-driven tethering. We have identified the role of phosphoinositide binding, and upstream lipid conversion, in regulating functional tethering by the exocyst complex. This lipid kinase-mediated mechanism provides a novel framework for understanding exocyst-dependent delivery of diverse cargo vesicles to the plasma membrane.

The direct observation that the exocyst holocomplex tethers two distinct PI(4,5)P_2_ membranes to each other requires the two lipid binding regions of the holocomplex to be in physical opposition. However, in current structures of the yeast exocyst complex (*38*, *46*), the localization of the lipid binding regions cannot be directly observed, owing to a large flexible linker connecting the PH domain of Sec3 (Exoc1) to the complex. The mammalian complex lacks this linker, likely resulting in a fixed and different location of the PH domain within the holocomplex. Indeed, the two PI(4,5)P_2_ binding regions of the yeast exocyst may be redundant, as vesicle tethering still occurs if either of them is deleted (*36*, *37*). In contrast, the mammalian exocyst requires both PI(4,5)P_2_ binding regions for the polarized trafficking of E-cadherin to cellular junctions (*33*), and the delivery of post-Golgi secretory vesicles to the plasma membrane (*35*). Our results highlight the importance of vesicular PI(4,5)P_2_ during polarized trafficking to the plasma membrane in a generalizable mechanism for exocyst-mediated tethering.

The generation of minor pools of PI(4,5)P_2_ on intracellular trafficking vesicles must be tightly regulated and places a requirement for the generation of PI(4,5)P_2_ downstream of endosome exit at the strictest of vicinity to the plasma membrane. Indeed, PI(4,5)P_2_ is present on internal membranes regulating distinct trafficking events (*65*). The most parsimonious model for these observations requires control over the localization of the lipid kinase, whose activity is directly linked to the activation of Arf6 (*15*). This model mechanistically implicates the activity of Arf6 on vesicles in both recycling (*49*) and potentially secretory pathways (*35*, *66*), yet its direct link to plasma membrane delivery at late steps has been mechanistically unclear. Our data uncover the direct role of Arf6-regulated PI(4,5)P_2_ generation and subsequent exocyst-mediated tethering in polarized trafficking.

Cargo proteins themselves may directly recruit PIP5K1C to trafficking vesicles (*33*), which would add an additional layer of complexity to this process. Indeed, further layers of regulatory complexity influence the composition of the exocyst holocomplex. The bivalent PI(4,5)P_2_ binding we have described for the exocyst is paralleled by Ral GTPase-mediated regulation, exerted through its independent binding to both Exoc2 and Exoc8 (*8*). Similarly, ULK1 and TBK1 regulate the exocyst through phosphorylation of Exoc7 and Exoc8 respectively (*8*, *67*, *68*). Our results highlight that these subunits are most capable of disassociation from the holocomplex, which while contrasting with some earlier suggestions (*69*), is supported by the hierarchal nature of the human exocyst (*40*, *43*).

Finally, the role of small GTPases in exocyst-dependent trafficking remains to be clarified. In particular, the activation and interconnections of Rab-family GTPases to Arf6 (*66*, *70*–*72*) are enticing in the context of the final steps of cargo carrier delivery to the plasma membrane. The novel mechanism we have uncovered for exocyst-mediated tethering provides a framework for linking the roles of the SNARE membrane fusion machinery, small GTPases, and the myriad of other exocyst binding partners to a cohesive model of polarized trafficking.

## Materials and Methods

### Plasmid construction

Exocyst subunit plasmids were a kind gift from Channing Der (Addgene 53755-53762) (*73*). Each was subcloned into FlexiBAC SF9 expression plasmids as GST-HRV3C, 6xHis-MBP-HRV3C, and 6xHis-HRV3C-mNeonGreen fusions, where HRV3C is a protease cleavage site, by standard molecular biology approaches (*45*). GFP-PIPK1 gamma 90 was a gift from Pietro De Camilli (Addgene plasmid # 22299) and subcloned with mScarlet-I into mammalian expression plasmids. Plasmids encoding PLCδ-PH and SidC were a kind gift from Evzen Boura and subcloned into bacterial expression plasmids. Gene encoding Arf6 was cloned into bacterial expression plasmids as a GST-fusion, and subcloned with mRuby3 into mammalian expression plasmids.

### Baculovirus production

Exocyst subunits and PIP5K1C were expressed in SF9 cells cultured in Insect-EXPRESS (Lonza) by baculovirus mediated expression. Baculovirus was produced using the FlexiBac methods (*45*). SF9 cells at 1×10^6^ cells/ml were co-transfected by the linearised Defbac viral backbone and the plasmids encoding for exocyst subunits or PIP5K1C using ESCORT-IV (Sigma). After 5 days, cells were checked for signs of viral transfection and 50-200 μl of supernatant of this P1 virus were used to infect 50 ml of insect cells to generate P2 virus. 5 days after infection, this P2 virus was again evaluated for virus production (LUNA-II; Logos Bio).

### Exocyst subunit interaction screen

For identification of exocyst subunit interactions, SF9 cells at 1×10^6^ SF9 cells/ml were grown in 24 well round bottom plates (Qiagen) and infected with baculovirus of the indicated single subunits. After 2 days of infection, the cells were pelleted at 500xg for 10 min, the supernatant discarded and the pellet frozen in liquid nitrogen. Upon use, the pellet was thawed and resuspended in 2 ml of standard buffer (20mM HEPES, 250mM NaCl, 0.5 mM TCEP), sonicated, and clarified at 4 °C. 25 μl of glutathione agarose beads (MRC-PPU Reagents and Services) were added and incubated for 1-2 h at 4 °C. Beads were isolated by centrifugation and washed three times in standard buffer using MobiSpin Column F (MoBiTec GmbH). The washed beads were resuspended in 50 μl 1x SDS loading buffer, boiled at 95 °C, and separated by SDS-PAGE. Gels were stained with Quick Coomassie Stain (Generon) and scanned.

### Protein Purifications

Exocyst complexes and PIP5K1C were expressed using the SF9 insect cell system (*45*). To express exocyst Subcomplex-1, cells were coinfected with N-terminal 6xHis-MBP tagged Exoc1 and untagged Exoc2. Separately, cells were coinfected with untagged Exoc3 and N-terminal 6xHis-mNeonGreen tagged Exoc4. Exoc1 was infected at 0.9 relative MOI relative to the other subunits to assure complex integration, with each virus having similar MOI. After two days of infection, the cultures were mixed, pelleted and frozen in liquid nitrogen.

To express exocyst Subcomplex-2, cells were coinfected with N-terminal 6xHis-mNeonGreen tagged Exoc6 and N-terminal 6xHis-MBP tagged Exoc5 at 0.8 relative MOI. Separately, untagged Exoc7 and untagged Exoc8 were coexpressed at 1.2 relative MOI. After two days of infection, the cultures were mixed, pelleted and frozen in liquid nitrogen.

To express exocyst holocomplex, cells were coinfected with untagged Exoc1 and untagged Exoc2, and separately with untagged Exoc3 and untagged Exoc4. Separately, cells were coinfected with N-terminal 6xHis-MBP tagged Exoc5 at 0.8 relative MOI, and N-terminal 6xHis-mNeonGreen tagged Exoc6. Finally, a final separate culture was infected with untagged Exoc7 and untagged Exoc8 at 1.2 relative MOI. After two days of infection, each of the 4 cultures was combined, pelleted and frozen in liquid nitrogen. In our hands, the MOI ratio of tagged subunit was of critical importance for generation of stoichiometric complexes.

To express PIP5K1C, cells were infected with N-terminal 6xHis-MBP tagged PIP5K1C for two days, pelleted and frozen in liquid nitrogen.

To purify the exocyst complexes, all pellets were thawed on ice and resuspended to 50ml final volume in standard buffer with the addition of protease inhibitors and Benzonase (Sigma). PIP5K1C was resuspended in buffer containing 20 mM HEPES, 400 mM NaCl, 0.5 mM TCEP, 50 mM Na2HPO4. Cell suspensions were lysed by Dounce homogenization followed by clarification through centrifugation. Cleared lysate was filtered through 0.45 μm membranes and bound to preequilibrated Amylose Fast Flow resin (MRC-PPU Reagents and Services). After washing, protein/proteins complexes were cleaved by addition of recombinant HRV-3C protease for at least 3 hours. Following cleavage and centrifugation, the supernatant was concentrated to using Vivaspin concentrators (Cytiva) and injected onto a 24 ml Superose 6 Increase 10/300 GL (or Superdex200 10/300 GL for Exoc1:mNeonGreen-Exoc4 and Exoc5:mNeonGreen-Exoc6:Exoc7) equilibrated in standard buffer. Peak fractions were analyzed by SDS-PAGE, pooled and concentrated and either used immediately in experiments, or aliquoted and frozen in liquid nitrogen and stored at −80 °C. Samples were sent to the mass spectrometry facility (Fingerprints, University of Dundee) for quantitative peptide analysis.

Small GTPases and lipid probes were expressed in BL21 bacterial cells using standard approaches. All constructs were grown in LB containing 1.75 w/vol % lactose and antibiotic at 37 °C to OD_600_=0.8, whereupon temperature was lowered to 18 °C for 10-12 h. Cells were pelleted resuspended in standard buffer, lysed, and clarified. Protein was purified by Ni affinity chromatography using 5 ml His-Trap HP column (Cytiva) followed by anion exchange on a 5ml Capto-Q column (Cytiva). Peak fractions were pooled and purified by size exclusion chromatography using a Superdex 200 16/60 pg column (Cytiva). All proteins were aliquoted, frozen in liquid nitrogen and stored at −80 °C.

### Negative stain electron microscopy

Carbon Film, 400 Mesh copper grids (Electron Microscopy Sciences) were glow discharged and exocyst holocomplex (1.9 μM) was applied for 10 minutes and blotted away with Whatman paper. Grids where washed twice with ddH2O before being stained with 0.75% uranyl formate for approximately 30s. Excess stain was removed by blotting, and samples were imaged using a JEOL 2200FS 200 kV with inline omega filter on a Gatan Ultrascan 4k 15 μm pixel camera at 30,000x magnification. Images were obtained at a defocus of −0.5-2.0 μm. Particles were manually selected using Boxer implemented in EMAN2 (*74*). Morphological model was generated in EMAN2 from 7052 particles over 75 micrographs to assess protein complex heterogeneity and dimensions.

### Lipids, membrane-coated beads, and lipid binding experiments

All lipids were obtained from Avanti Polar Lipids unless otherwise stated. Liposomes containing either PI(4)P (Echelon Biosciences) or PI(4,5)P_2_ (Echelon Biosciences) were produced by mixing 95 mol % 1-palmitoyl-2-oleoyl-glycero-3-phosphocholine (POPC) with 5 mol % of the respective phosphatidylinositol together with 0.1% Atto647N-DOPE (ATTO-TEC) or Rhodamine-DOPE. The mixtures were evaporated under nitrogen flow and dried overnight in a vacuum. Dried lipids were resuspended in buffer containing 20 mM HEPES, 150 mM NaCl, and 0.5 mM TCEP at 37 °C and subjected to 6 cycles of freeze thawing in liquid nitrogen. Where samples contained 0.5% Tris-NTA-DODA (*62*) for Arf6 conjugation, 1 mM NiCl2 was added during the resuspension step. Liposomes were extruded to 100 nm and aliquoted, frozen in liquid nitrogen and stored at −20 °C.

Membrane-coated beads (*51*, *52*) were generated by mixing liposomes and 10 μm silica beads (Whitehouse Scientific) in 200 mM NaCl. Beads were washed twice with 20 mM HEPES and resuspended in buffer containing 20 mM HEPES, 150 mM NaCl, and 0.5mM TCEP. Samples were kept on a rotator until use within 2 hours of generation.

To assess lipid binding, beads were added to uncoated μ-Slide 8 well chambers (Ibidi) and protein added and mixed by pipetting at the indicated final concentrations. Lipid binding kinetics were allowed to equilibrate for 30 min at room temperature before imaging at the transverse plane.

Confocal images were acquired using a Leica SP8 Confocal Microscope with a Leica HC PL APO CS2 63x/1.40 Oil objective at 0.75 base zoom with 1024×1024 pixels scan. Data was analysed using a custom written ImageJ script. The signal from the Atto647N-DOPE or Rhodamine-DOPE was used to segment each bead individually, thereby creating a mask for the circumference for bead. The mask was used in the GFP/mNeonGreen channel to measure the recruitment of the fluorescent probes to each bead. Statistics shown are single-site model, probability by Aoike Information Criterion <0.01%.

### Phosphoinositide conversion assays

Membrane-coated beads containing 0.1% Atto647N-DOPE (ATTO-TEC), 5% PI(4)P and 10% phosphatidyl serine (PS) in 84.9% POPC were prepared as described above and resuspended in 100 μl kinase buffer (20 mM HEPES (pH 7.0), 150 mM NaCl, 5 mM MgCl2, 0.5 mM EGTA, 200 μg/mL β-casein, 20 mM BME, and 20 mM glucose) in the presence or absence of 1 mM ATP. Samples were applied to an imaging chamber and protein was added to a final concentration of 100 nM. Immediately before acquisition, reconstituted PIP5K1C was added to the beads at a final concentration of 150 nM. The reaction was monitored at 0.2 Hz using a Leica SP8 Confocal Microscope with a Leica HC PL APO CS2 63x/1.40 Oil objective at 0.75 base zoom with 1024×1024 pixel scan.

For membrane tethering assays, liposomes containing 5% PI(4,5)P_2_ were added to the beads together with PIP5K1C for 30 min while rotation at RT before imaging.

For Arf6 mediated phosphoinositide conversion, 1 μM purified 6xHis-Arf6-T27N/Q67L was added to membrane-coated beads containing 0.5% Tris-NTA-DODA (*62*), 5% PI4P, 10% PS in 84.5% POPC doped with either 0.1% NBD-DPPE or Atto647N-DOPE (ATTO-TEC) for 30 min and washed three times. At these conditions Arf6 binding to the beads were saturated. Beads were added to an Ibidi imaging slide and purified PLCδ-PH domain with N-terminal RFP tag were added to a final concentration of 100 nM. Immediately before imaging, purified PIP5K1C was added to the beads at a final concentration of 12.5 nM. Recruitment of PLCδ to either of the two membrane-coated bead populations was determined using ImageJ by segmentation of either the NBD or Atto647N signal as mask.

### Cryo-electron tomography

Exocyst holocomplex was mixed with liposomes composed of 95 mol% POPC and 5 mol% PI(4,5)P_2_ and protein A coated 5 nm gold fiducials (Electron Microscopy Sciences) for 3 minutes and to a final protein concentration of 1 μM. Samples were applied to glow-discharged holey carbon film grids (Quantifoil) and plunge frozen into liquid ethane using a Vitrobot (ThermoFisher). Tilt series were collected on a JEOL CRYO ARM 300 (Jeol Ltd) electron microscope operated at 300 keV and equipped with in-column omega energy filter and DE-64 direct electron detector (Direct Electron, USA) operated in electron counting mode. Data acquisition was controlled by SerialEM (*75*) using the dose-symmetric scheme (*76*) using an increment of 1° and a group size of 3° with a range of −60° to +60°. The nominal magnification was 30,000x, giving a calibrated pixel size of 2.35 Å/px. The defocus was set to −6 μm and the dose was 1.84 e/Å2/tilt giving a total dose of 223 e/Å2 over a complete tilt series. The energy filter was set to a slit width of 30 eV. Images were collected as movies of 6 frames per tilt. Movies were motion corrected, gain corrected, assembled into tilt series and dose weighted using the IMOD program alignframes (*75*). Tomograms were reconstructed by back projection using the IMOD program ETOMO with 5 nm gold particles as fiducials. Liposomes were segmented by hand throughout the tilt series in 3dmod (*77*) and the distance of closest approach was determined. Proteinaceous material that was clearly associated with liposomes was segmented using the AutoThreshold function.

### Live cell Airyscan imaging

NMuMG cells (ATCC) and NMuMG genome-modified cells (*43*) were grown in DMEM containing 4.5 μg/ml Glucose with Pyruvate, supplemented with 10% FBS and PenStrep. Cells were seeded onto Ibidi glass bottom dishes and transfected using NMuMG-optimized GeneJet transfection reagent (ThermoFisher) according to the manufacturer’s instructions. 24-48 h after transfection cells were imaged in an environmental control chamber at 37 C with 5% CO_2_ using a Zeiss LSM880 Airyscan microscope with a 60x oil objective NA1.4. Overlap between the different channels was quantified in ImageJ using automated segmentation tool Squassh (*78*). Signal overlap was determined in Squassh (*78*) and plotted as a SuperPlot (*79*) with bold symbols representing the mean of an individual biological repeat, and the background symbols representing the data from individual cells.

### Phosphoinositide staining in cells

Staining protocols for phosphoinositides using purified SidC-GFP and GFP-PLCδ-PH were adopted from Hammond et al 2019. For staining with purified SidC-GFP, cells were fixed with 37 °C prewarmed PBS with 4% PFA for 20 min, followed by quenching in 50 mM NH4Cl for 20 min. Fixed cells were permeabilised with 20 μM digitonin in 1X PIPES buffer for 5 min at room temperature and washed. Fixed and permeabilised cells were blocked with 5% goat serum in 50 mM NH4Cl containing PIPES buffer in the presence of 1 μM purified SidC-GFP for 1 h at room temperature. Cells were washed in PIPES buffer and stained with mouse anti-GM130 (BD Biosciences 610822) for 1 h, washed and stained with secondary antibodies coupled to Alexa 647 for 45 min, washed and postfix in PBS with 2% PFA for 10 min. Samples were washed in 50 mM NH4Cl, mounted using Prolong Gold (ThermoFisher) and imaged using an LSM880 Airyscan microscope with a 60x, NA1.4 oil objective.

For staining with purified GFP-PLCδ-PH, cells were fixed with prewarmed 37C 4% PFA in PBS for 20min, followed by quenching in 50 mM NH4Cl for 20 min. From now on all steps were carried out on ice and with all solutions chilled to 4 °C. Fixed cells were permeabilised with 0.5% saponin in PIPES buffer with 5% goat serum and 2 μM purified GFP-PLCδ-PH for 1 h. Fixed and permeabilised cells were washed with PIPES buffer and stained with mouse anti-e-cadherin (Clone 36/E; BD Bioscience) in 5% goat serum and 0.1% saponin containing PIPES buffer for 1 h washed and stained with secondary antibodies coupled to Alexa 647 for 45 min. Ice cold PBS with 2% PFA was added for 10 min before transferring the slides to room temperature and letting them adjust for another 10 min. Samples were rinsed in 50 mM NH4Cl, mounted, and imaged as above described.

Signal overlap was determined in Squassh (*78*) and plotted as a SuperPlot (*79*) with bold symbols representing the mean of an individual biological repeat, and the background symbols representing the data from individual cells.

### Arf6 - PIP5K pulldown

BL21 cells transformed with GST alone or GST-Arf6-Q67L/T27N were grown in LB containing 1.75% lactose and antibiotic at 37 °C to OD_600_=0.8, whereupon temperature was lowered to 18 °C for 10-12 h. Cells were pelleted, resuspended in standard buffer, lysed, and clarified. Sample was then filtered through a 0.22 μm filter and bound to GSH agarose beads (MRC-PPU Reagents and Services) for 1 h at 4 °C. Beads were washed, and protein was eluted with 20 μM glutathione. Eluted protein was further purified by size exclusion chromatography using a Superdex 200 16/60 pg column. Peak fractions were pooled and snap frozen in liquid nitrogen. Equal amounts of purified protein were added to 50 μl GSH agarose for 1 h at 4 °C, washed and resuspended in standard buffer. Purified PIP5K1C was added to a final concentration of 400 nM and incubated at 4 °C under constant rotation for 1 h. Beads were washed, resuspended in SDS-loading buffer and analysed by SDS page, followed by Coomassie staining.

## Acknowledgments

We are very grateful to Ian Macara and Mukhtar Ahmed for sharing their cell lines. We thank Hayley Shaw, Aliona Bogdanova, Evzen Boura lab, and Jakub Piehler lab for reagents. We acknowledge Ramasubramaniam Sundaramoorthy for EM assistance. We thank Jens Januschke, Jason King, and Kees Weijer for their comments on the manuscript. We gratefully acknowledge the Dundee Imaging Facility, University of Dundee for their support in this work. We acknowledge the Scottish Centre for Macromolecular Imaging (SCMI) and James Streetley for assistance with cryo-EM experiments and access to instrumentation, funded by the MRC (MC_PC_17135) and SFC (H17007). We would like to acknowledge the FingerPrints Proteomics Facility at the University of Dundee, which is supported by the ‘Wellcome Trust Technology Platform’ award [097945/B/11/Z].

## Funding

This work was supported by the

Wellcome Trust grant 211193/Z/18/Z to DHM;

Royal Society RGS\R2\180284 to DHM.

## Author contributions

The research reported emerged from discussions between both authors, and all authors contributed to the writing of the manuscript.

## Supplemental material

**Fig. S1.**
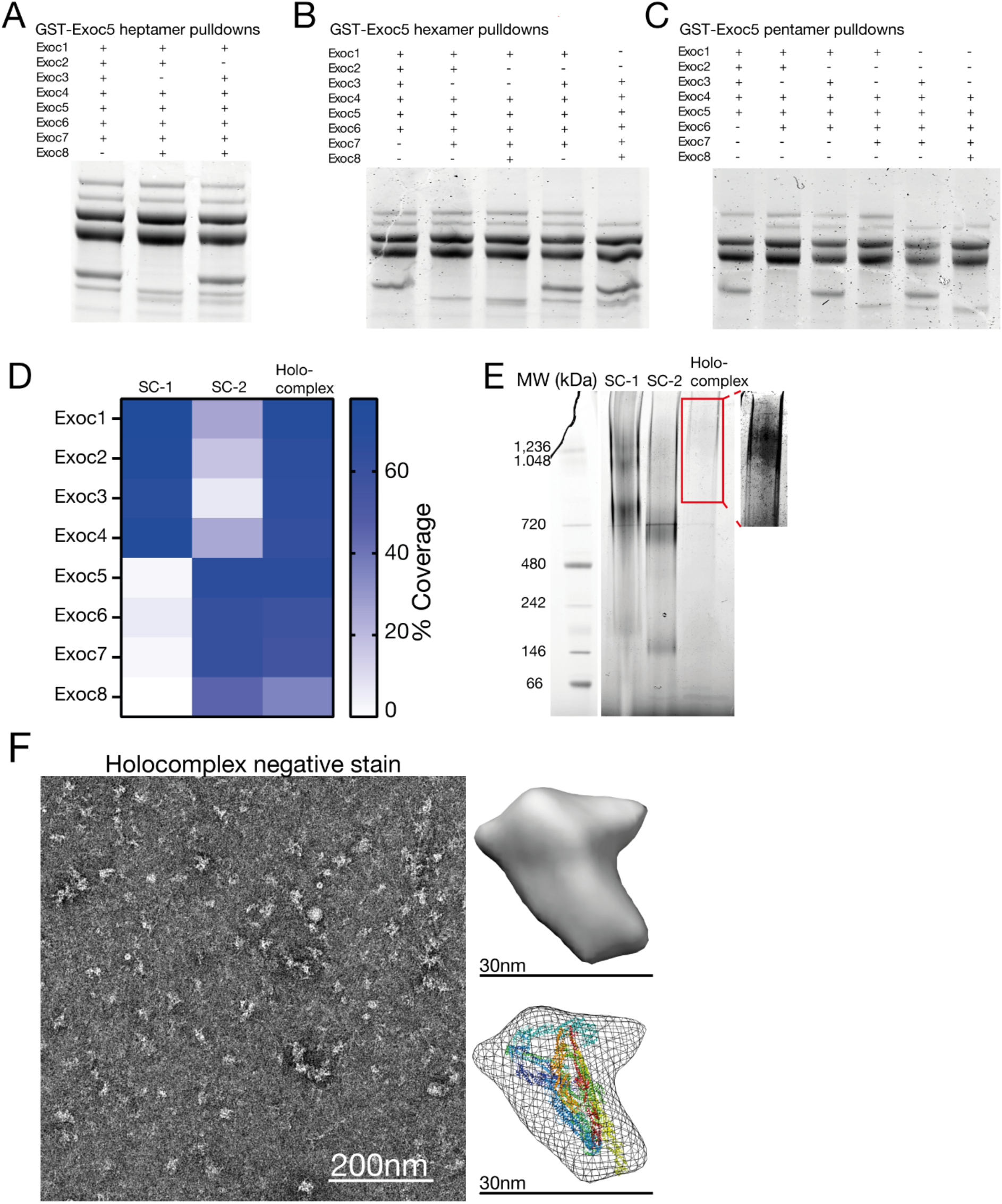
Exocyst connectivity and holocomplex verification and validation. **A-C**, To confirm higher order exocyst complex connectivity, exocyst subunits were separately expressed in SF9 insect cells and mixed upon lysis. All possible combinations for exocyst hepta-, hexa-, and pentamers from pulldowns using Exoc5 tagged with GST and analysed by SDS-PAGE and Coomassie staining. **D**, Heatmap of protein coverage for mass spectrometry analysis of purified Subcomplex-1, −2 and holocomplex. **E**, Native gel electrophoresis of purified Subcomplex-1, −2 and holocomplex. Insert is shown in enhanced contrast to visualise faint bands from holocomplex. F, Exocyst holocomplex was analysed by negative stain and a low-resolution model was generated. The cryo-EM structure of the yeast exocyst (ribbons) was placed into the outline (mesh) of this low-resolution shell for visual comparison.

**Fig. S2.**
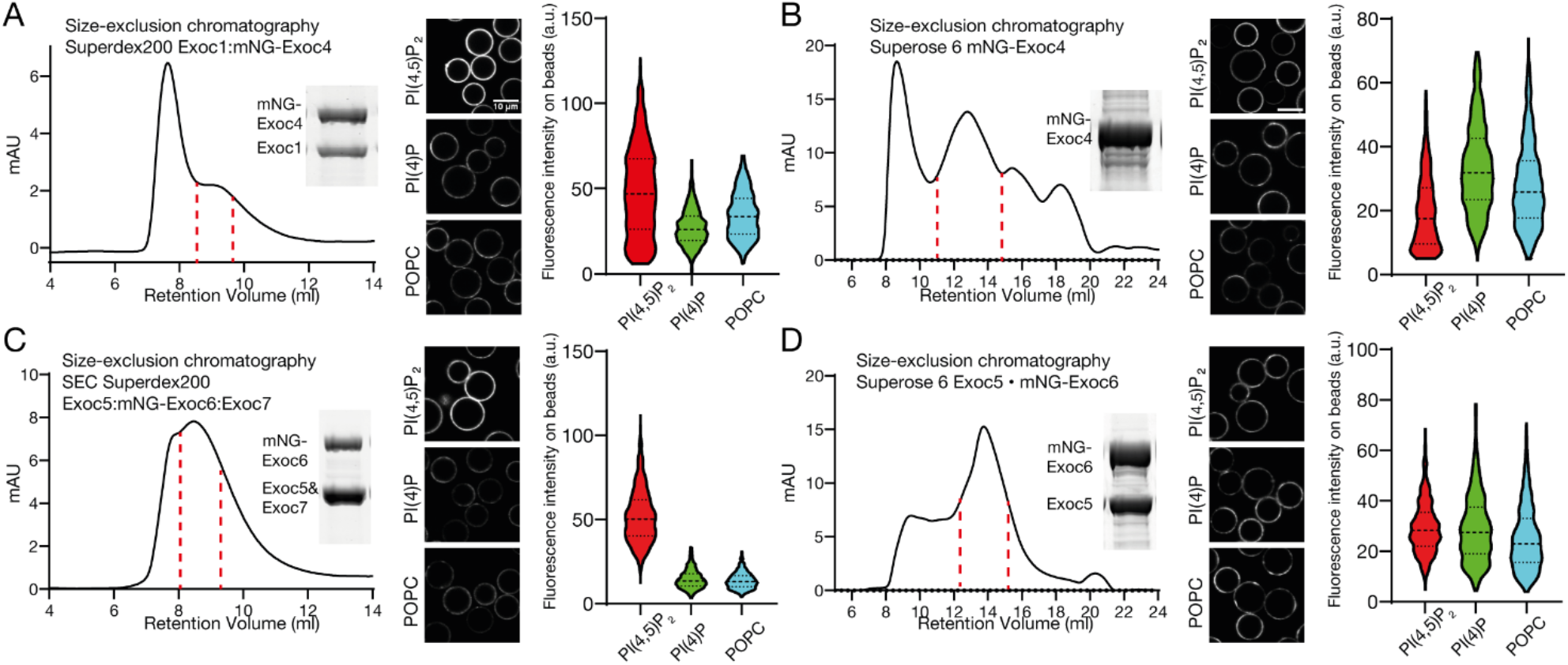
Purification and phosphoinositide binding of exocyst subcomplexes. **A**, Size-exclusion chromatography of a dimer of Exoc1 with mNeonGreen (mNG)-tagged Exoc4. **B**, Exoc1:4 dimer was added to membrane-coated beads. These were formed from a lipid composition of 84.9% 1-palmitoyl-2-oleoyl-glycero-3-phosphocholine (POPC), 10% phosphatidylserine and 5% phosphoinositide or POPC, doped with 0.1% rhodamine-DPPE. Individual beads were segmented, and mean fluorescent intensity was quantified in ImageJ. n=926-1589 beads. **C**, Size-exclusion chromatography of mNeonGreen (mNG) tagged Exoc4. **D**, Exoc4-mNG was added to membrane-coated beads as in B. n=745-881 beads. **E**, Size-exclusion chromatography of trimer of Exoc5:6:7 with mNeonGreen (mNG)-tagged Exoc6. **F**, Exoc5:6:7 timer was added to membrane-coated beads as in B. n=1032-1239 beads. **G**, Size-exclusion chromatography of dimer of Exoc5:6 with mNeonGreen (mNG) tagged Exoc6. **H**, Exoc5-6 dimer was added to membrane-coated beads as in B. n=1025-1178 beads.

**Fig. S3.**
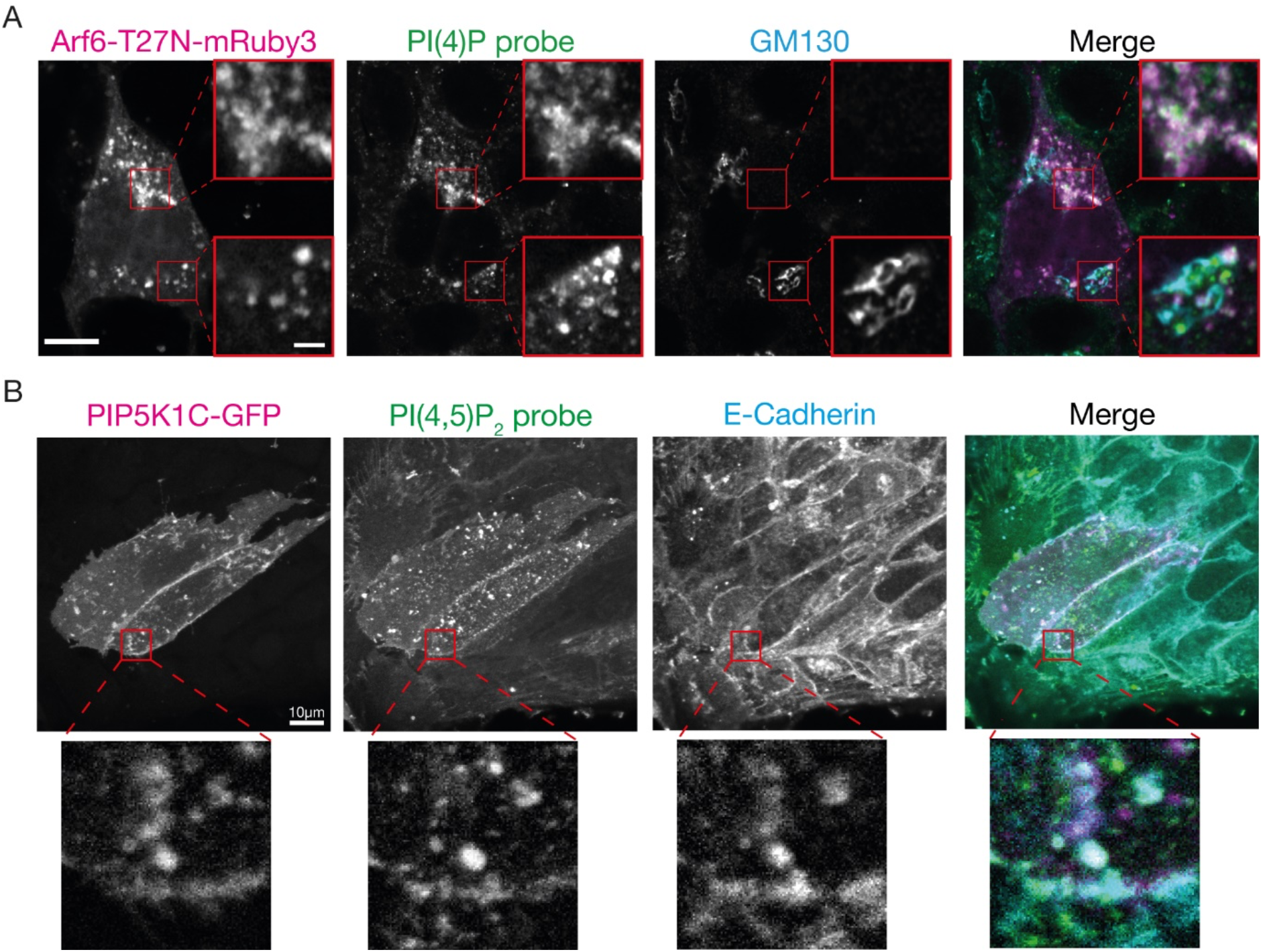
PI(4)P and PI(4,5)P_2_ staining of fixed and permeabilized NMuMG cells. **A**, NMuMG cells were transfected with Arf6-T27N tagged with mRuby3 and stained for PI(4)P using recombinant SidC fused to GFP and conventional antibody staining against GM130. **B**, NMuMG cells were transfected with PIP5K1C tagged with EGFP and stained for PI(4,5)P_2_ using recombinant PLCδ-PH fused to RFP, and conventional antibody staining against e-cadherin. Scale bar, 10 μm, inset 1 μm.

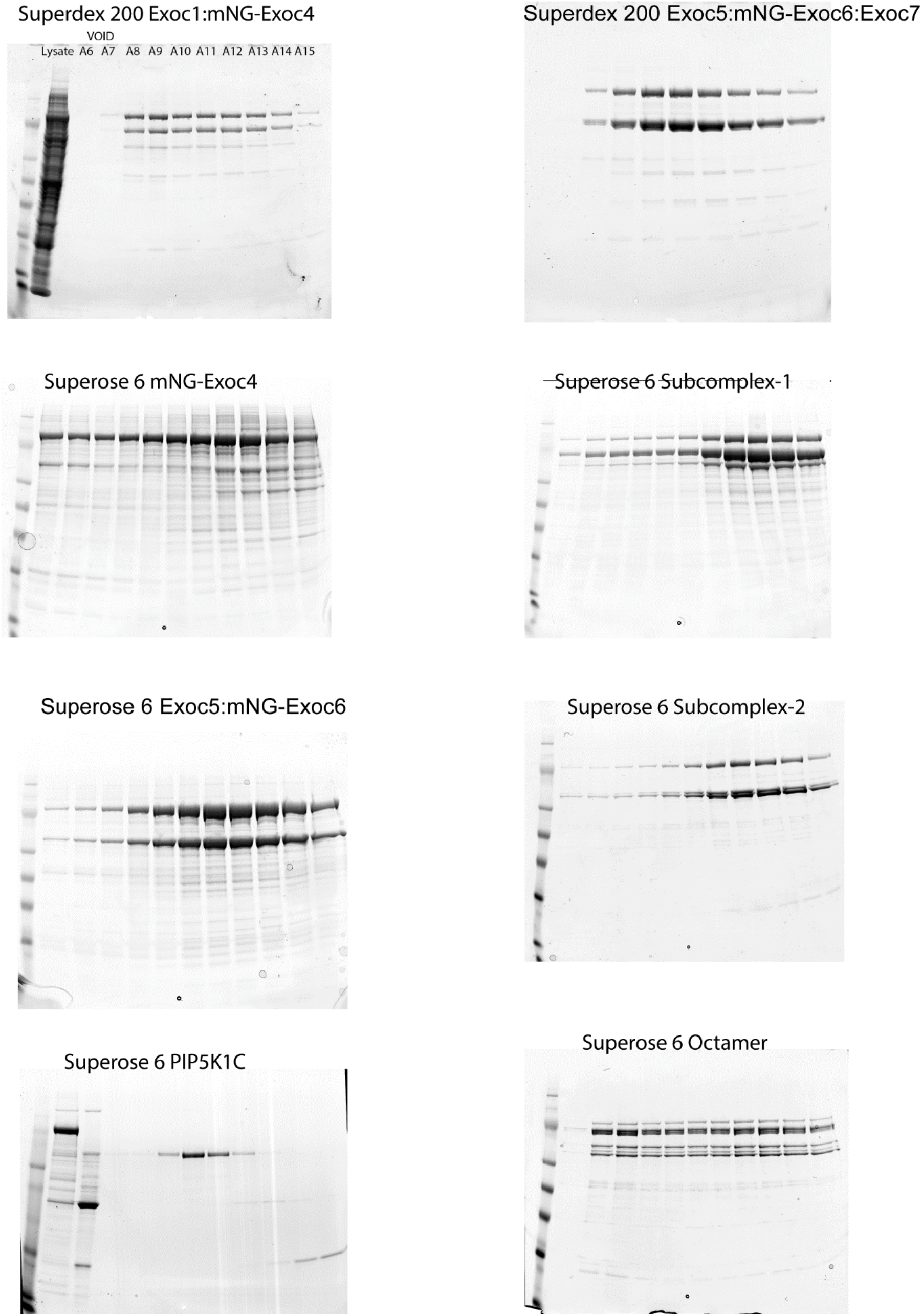

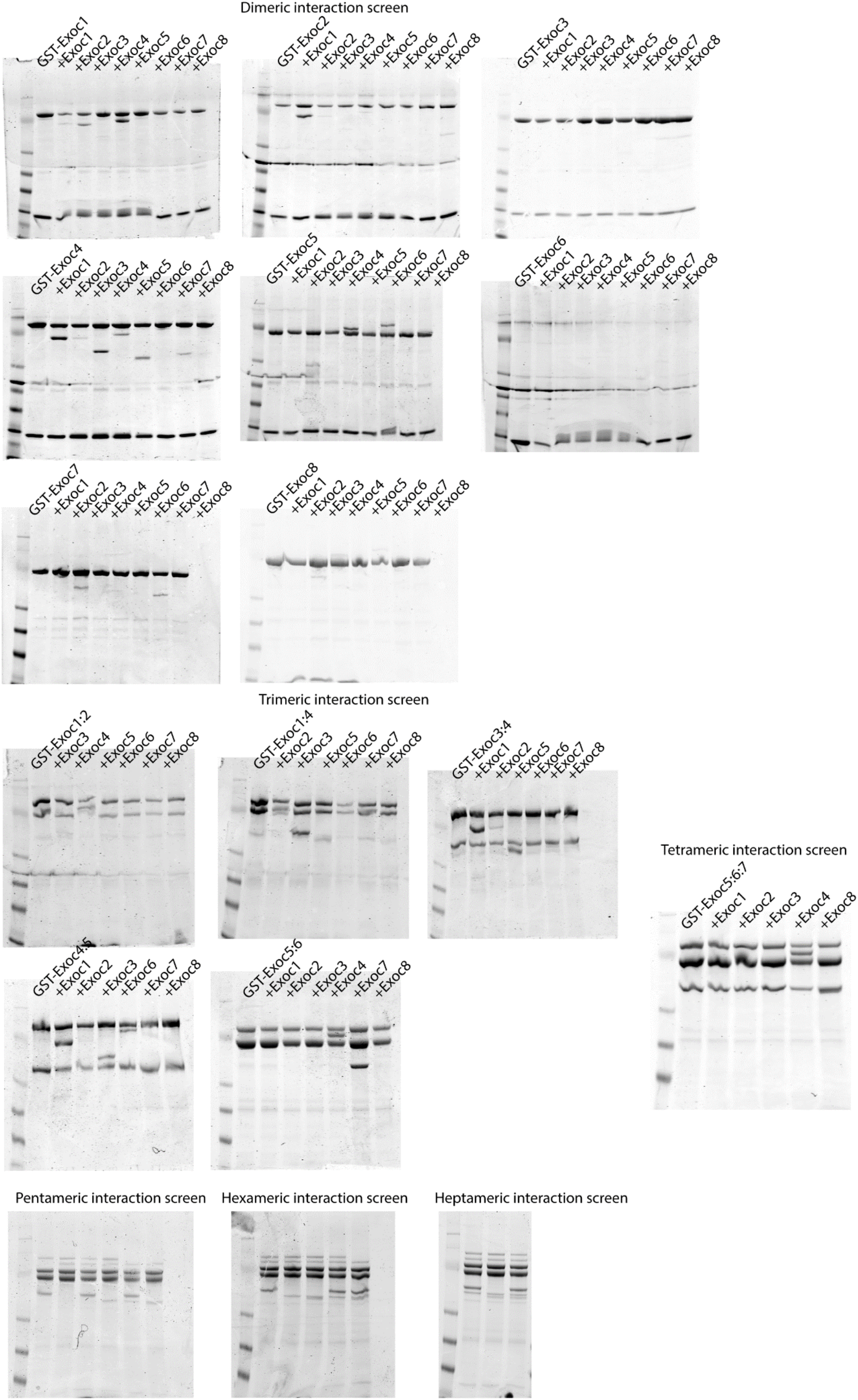

## References

1. D. M. Bryant, K. E. Mostov, From cells to organs: building polarized tissue. Nat Rev Mol Cell Biol 9, 887–901 (2008).

2. B. Wu, W. Guo, The Exocyst at a Glance. J Cell Sci 128, 2957–2964 (2015).

3. T. Lecuit, F. Pilot, Developmental control of cell morphogenesis: a focus on membrane growth. Nat Cell Biol 5, 103–108 (2003).

4. R. Wang, C. R. Simoneau, J. Kulsuptrakul, M. Bouhaddou, K. A. Travisano, J. M. Hayashi, J. Carlson-Stevermer, J. R. Zengel, C. M. Richards, P. Fozouni, J. Oki, L. Rodriguez, B. Joehnk, K. Walcott, K. Holden, A. Sil, J. E. Carette, N. J. Krogan, M. Ott, A. S. Puschnik, Genetic Screens Identify Host Factors for SARS-CoV-2 and Common Cold Coronaviruses. Cell 184, 106–119 e114 (2021).

5. P. Novick, C. Field, R. Schekman, Identification of 23 complementation groups required for post-translational events in the yeast secretory pathway. Cell 21, 205–215 (1980).

6. D. R. TerBush, T. Maurice, D. Roth, P. Novick, The Exocyst is a multiprotein complex required for exocytosis in Saccharomyces cerevisiae. Embo j 15, 6483–6494 (1996).

7. S. C. Hsu, C. D. Hazuka, D. L. Foletti, R. H. Scheller, Targeting vesicles to specific sites on the plasma membrane: the role of the sec6/8 complex. Trends Cell Biol 9, 150–153 (1999).

8. S. Moskalenko, C. Tong, C. Rosse, G. Mirey, E. Formstecher, L. Daviet, J. Camonis, M. A. White, Ral GTPases regulate exocyst assembly through dual subunit interactions. J Biol Chem 278, 51743–51748 (2003).

9. M. L. Dubuke, S. Maniatis, S. A. Shaffer, M. Munson, The Exocyst Subunit Sec6 Interacts with Assembled Exocytic SNARE Complexes. J Biol Chem 290, 28245–28256 (2015).

10. P. Yue, Y. Zhang, K. Mei, S. Wang, J. Lesigang, Y. Zhu, G. Dong, W. Guo, Sec3 promotes the initial binary t-SNARE complex assembly and membrane fusion. Nat Commun 8, 14236 (2017).

11. C. Rosse, E. Formstecher, K. Boeckeler, Y. Zhao, J. Kremerskothen, M. D. White, J. H. Camonis, P. J. Parker, An aPKC-exocyst complex controls paxillin phosphorylation and migration through localised JNK1 activation. PLoS Biol 7, e1000235 (2009).

12. G. Rossi, D. Lepore, L. Kenner, A. B. Czuchra, M. Plooster, A. Frost, M. Munson, P. Brennwald, Exocyst structural changes associated with activation of tethering downstream of Rho/Cdc42 GTPases. J Cell Biol 219, (2020).

13. S. M. Ahmed, I. G. Macara, The Par3 polarity protein is an exocyst receptor essential for mammary cell survival. Nat Commun 8, 14867 (2017).

14. J. E. Burke, A. J. Inglis, O. Perisic, G. R. Masson, S. H. McLaughlin, F. Rutaganira, K. M. Shokat, R. L. Williams, Structures of PI4KIIIbeta complexes show simultaneous recruitment of Rab11 and its effectors. Science 344, 1035–1038 (2014).

15. Y. Aikawa, T. F. Martin, ARF6 regulates a plasma membrane pool of phosphatidylinositol(4,5)bisphosphate required for regulated exocytosis. J Cell Biol 162, 647–659 (2003).

16. A. Honda, M. Nogami, T. Yokozeki, M. Yamazaki, H. Nakamura, H. Watanabe, K. Kawamoto, K. Nakayama, A. J. Morris, M. A. Frohman, Y. Kanaho, Phosphatidylinositol 4-phosphate 5-kinase alpha is a downstream effector of the small G protein ARF6 in membrane ruffle formation. Cell 99, 521–532 (1999).

17. C. Marquer, H. Tian, J. Yi, J. Bastien, C. Dall’Armi, Y. Yang-Klingler, B. Zhou, R. B. Chan, G. Di Paolo, Arf6 controls retromer traffic and intracellular cholesterol distribution via a phosphoinositide-based mechanism. Nat Commun 7, 11919 (2016).

18. A. Shi, O. Liu, S. Koenig, R. Banerjee, C. C. Chen, S. Eimer, B. D. Grant, RAB-10-GTPase-mediated regulation of endosomal phosphatidylinositol-4,5-bisphosphate. Proc Natl Acad Sci U S A 109, E2306–2315 (2012).

19. M. Prigent, T. Dubois, G. Raposo, V. Derrien, D. Tenza, C. Rosse, J. Camonis, P. Chavrier, ARF6 controls post-endocytic recycling through its downstream exocyst complex effector. J Cell Biol 163, 1111–1121 (2003).

20. V. L. Koumandou, J. B. Dacks, R. M. Coulson, M. C. Field, Control systems for membrane fusion in the ancestral eukaryote; evolution of tethering complexes and SM proteins. BMC Evol Biol 7, 29 (2007).

21. L. Yakir-Tamang, J. E. Gerst, A phosphatidylinositol-transfer protein and phosphatidylinositol-4-phosphate 5-kinase control Cdc42 to regulate the actin cytoskeleton and secretory pathway in yeast. Mol Biol Cell 20, 3583–3597 (2009).

22. L. S. Garrenton, C. J. Stefan, M. A. McMurray, S. D. Emr, J. Thorner, Pheromone-induced anisotropy in yeast plasma membrane phosphatidylinositol-4,5-bisphosphate distribution is required for MAPK signaling. Proc Natl Acad Sci U S A 107, 11805–11810 (2010).

23. E. Mizuno-Yamasaki, M. Medkova, J. Coleman, P. Novick, Phosphatidylinositol 4-phosphate controls both membrane recruitment and a regulatory switch of the Rab GEF Sec2p. Dev Cell 18, 828–840 (2010).

24. B. He, W. Guo, The exocyst complex in polarized exocytosis. Curr Opin Cell Biol 21, 537–542 (2009).

25. N. Thapa, Y. Sun, M. Schramp, S. Choi, K. Ling, R. A. Anderson, Phosphoinositide signaling regulates the exocyst complex and polarized integrin trafficking in directionally migrating cells. Dev Cell 22, 116–130 (2012).

26. G. Di Paolo, P. De Camilli, Phosphoinositides in cell regulation and membrane dynamics. Nature 443, 651–657 (2006).

27. A. Wallroth, V. Haucke, Phosphoinositide conversion in endocytosis and the endolysosomal system. J Biol Chem 293, 1526–1535 (2018).

28. D. H. Murray, L. K. Tamm, Clustering of syntaxin-1A in model membranes is modulated by phosphatidylinositol 4,5-bisphosphate and cholesterol. Biochemistry 48, 4617–4625 (2009).

29. S. J. An, F. Rivera-Molina, A. Anneken, Z. Xi, B. McNellis, V. I. Polejaev, D. Toomre, An active tethering mechanism controls the fate of vesicles. Nat Commun 12, 5434 (2021).

30. K. Ketel, M. Krauss, A. S. Nicot, D. Puchkov, M. Wieffer, R. Muller, D. Subramanian, C. Schultz, J. Laporte, V. Haucke, A phosphoinositide conversion mechanism for exit from endosomes. Nature 529, 408–412 (2016).

31. C. C. Campa, J. P. Margaria, A. Derle, M. Del Giudice, M. C. De Santis, L. Gozzelino, F. Copperi, C. Bosia, E. Hirsch, Rab11 activity and PtdIns(3)P turnover removes recycling cargo from endosomes. Nat Chem Biol 14, 801–810 (2018).

32. T. Farmer, S. Xie, N. Naslavsky, J. Stockli, D. E. James, S. Caplan, Defining the protein and lipid constituents of tubular recycling endosomes. J Biol Chem 296, 100190 (2021).

33. X. Xiong, Q. Xu, Y. Huang, R. D. Singh, R. Anderson, E. Leof, J. Hu, K. Ling, An association between type Igamma PI4P 5-kinase and Exo70 directs E-cadherin clustering and epithelial polarization. Mol Biol Cell 23, 87–98 (2012).

34. J. Jouette, A. Guichet, S. B. Claret, Dynein-mediated transport and membrane trafficking control PAR3 polarised distribution. Elife 8, (2019).

35. J. Liu, X. Zuo, P. Yue, W. Guo, Phosphatidylinositol 4,5-bisphosphate mediates the targeting of the exocyst to the plasma membrane for exocytosis in mammalian cells. Mol Biol Cell 18, 4483–4492 (2007).

36. B. He, F. Xi, X. Zhang, J. Zhang, W. Guo, Exo70 interacts with phospholipids and mediates the targeting of the exocyst to the plasma membrane. EMBO J 26, 4053–4065 (2007).

37. X. Zhang, K. Orlando, B. He, F. Xi, J. Zhang, A. Zajac, W. Guo, Membrane association and functional regulation of Sec3 by phospholipids and Cdc42. J Cell Biol 180, 145–158 (2008).

38. K. Mei, Y. Li, S. Wang, G. Shao, J. Wang, Y. Ding, G. Luo, P. Yue, J. J. Liu, X. Wang, M. Q. Dong, H. W. Wang, W. Guo, Cryo-EM structure of the exocyst complex. Nat Struct Mol Biol 25, 139–146 (2018).

39. M. R. Heider, M. Gu, C. M. Duffy, A. M. Mirza, L. L. Marcotte, A. C. Walls, N. Farrall, Z. Hakhverdyan, M. C. Field, M. P. Rout, A. Frost, M. Munson, Subunit connectivity, assembly determinants and architecture of the yeast exocyst complex. Nat Struct Mol Biol 23, 59–66 (2016).

40. Y. Katoh, S. Nozaki, D. Hartanto, R. Miyano, K. Nakayama, Architectures of multisubunit complexes revealed by a visible immunoprecipitation assay using fluorescent fusion proteins. J Cell Sci 128, 2351–2362 (2015).

41. D. Fasshauer, R. B. Sutton, A. T. Brunger, R. Jahn, Conserved structural features of the synaptic fusion complex: SNARE proteins reclassified as Q- and R-SNAREs. Proc Natl Acad Sci U S A 95, 15781–15786 (1998).

42. A. N. Lupas, J. Bassler, Coiled Coils - A Model System for the 21st Century. Trends Biochem Sci 42, 130–140 (2017).

43. S. M. Ahmed, H. Nishida-Fukuda, Y. Li, W. H. McDonald, C. C. Gradinaru, I. G. Macara, Exocyst dynamics during vesicle tethering and fusion. Nat Commun 9, 5140 (2018).

44. F. O. Bendezu, V. Vincenzetti, S. G. Martin, Fission yeast Sec3 and Exo70 are transported on actin cables and localize the exocyst complex to cell poles. PLoS One 7, e40248 (2012).

45. R. P. Lemaitre, A. Bogdanova, B. Borgonovo, J. B. Woodruff, D. N. Drechsel, FlexiBAC: a versatile, open-source baculovirus vector system for protein expression, secretion, and proteolytic processing. BMC Biotechnol 19, 20 (2019).

46. S. J. Ganesan, M. J. Feyder, I. E. Chemmama, F. Fang, M. P. Rout, B. T. Chait, Y. Shi, M. Munson, A. Sali, Integrative structure and function of the yeast exocyst complex. Protein Sci 29, 1486–1501 (2020).

47. A. Picco, I. Irastorza-Azcarate, T. Specht, D. Boke, I. Pazos, A. S. Rivier-Cordey, D. P. Devos, M. Kaksonen, O. Gallego, The In Vivo Architecture of the Exocyst Provides Structural Basis for Exocytosis. Cell 168, 400–412 e418 (2017).

48. Y. Zhai, D. Zhang, L. Yu, F. Sun, F. Sun, SmartBac, a new baculovirus system for large protein complex production. J Struct Biol X 1, 100003 (2019).

49. M. Desclozeaux, J. Venturato, F. G. Wylie, J. G. Kay, S. R. Joseph, H. T. Le, J. L. Stow, Active Rab11 and functional recycling endosome are required for E-cadherin trafficking and lumen formation during epithelial morphogenesis. Am J Physiol Cell Physiol 295, C545–556 (2008).

50. K. Simons, A. Wandinger-Ness, Polarized sorting in epithelia. Cell 62, 207–210 (1990).

51. D. H. Murray, M. Jahnel, J. Lauer, M. J. Avellaneda, N. Brouilly, A. Cezanne, H. Morales-Navarrete, E. D. Perini, C. Ferguson, A. N. Lupas, Y. Kalaidzidis, R. G. Parton, S. W. Grill, M. Zerial, An endosomal tether undergoes an entropic collapse to bring vesicles together. Nature 537, 107–111 (2016).

52. T. J. Pucadyil, S. L. Schmid, Supported bilayers with excess membrane reservoir: a template for reconstituting membrane budding and fission. Biophys J 99, 517–525 (2010).

53. S. D. Hansen, W. Y. C. Huang, Y. K. Lee, P. Bieling, S. M. Christensen, J. T. Groves, Stochastic geometry sensing and polarization in a lipid kinase-phosphatase competitive reaction. Proc Natl Acad Sci U S A 116, 15013–15022 (2019).

54. V. L. Kolossov, M. Sivaguru, J. Huff, K. Luby, K. Kanakaraju, H. R. Gaskins, Airyscan super-resolution microscopy of mitochondrial morphology and dynamics in living tumor cells. Microsc Res Tech 81, 115–128 (2018).

55. G. R. Hammond, T. Balla, Polyphosphoinositide binding domains: Key to inositol lipid biology. Biochim Biophys Acta 1851, 746–758 (2015).

56. M. Klima, D. J. Toth, R. Hexnerova, A. Baumlova, D. Chalupska, J. Tykvart, L. Rezabkova, N. Sengupta, P. Man, A. Dubankova, J. Humpolickova, R. Nencka, V. Veverka, T. Balla, E. Boura, Structural insights and in vitro reconstitution of membrane targeting and activation of human PI4KB by the ACBD3 protein. Sci Rep 6, 23641 (2016).

57. M. A. Lemmon, K. M. Ferguson, R. O’Brien, P. B. Sigler, J. Schlessinger, Specific and high-affinity binding of inositol phosphates to an isolated pleckstrin homology domain. Proc Natl Acad Sci U S A 92, 10472–10476 (1995).

58. C. D’Souza-Schorey, G. Li, M. I. Colombo, P. D. Stahl, A regulatory role for ARF6 in receptor-mediated endocytosis. Science 267, 1175–1178 (1995).

59. H. Radhakrishna, O. Al-Awar, Z. Khachikian, J. G. Donaldson, ARF6 requirement for Rac ruffling suggests a role for membrane trafficking in cortical actin rearrangements. J Cell Sci 112 (Pt 6), 855–866 (1999).

60. F. D. Brown, A. L. Rozelle, H. L. Yin, T. Balla, J. G. Donaldson, Phosphatidylinositol 4,5-bisphosphate and Arf6-regulated membrane traffic. J Cell Biol 154, 1007–1017 (2001).

61. M. Krauss, M. Kinuta, M. R. Wenk, P. De Camilli, K. Takei, V. Haucke, ARF6 stimulates clathrin/AP-2 recruitment to synaptic membranes by activating phosphatidylinositol phosphate kinase type Igamma. J Cell Biol 162, 113–124 (2003).

62. O. Beutel, J. Nikolaus, O. Birkholz, C. You, T. Schmidt, A. Herrmann, J. Piehler, High-fidelity protein targeting into membrane lipid microdomains in living cells. Angew Chem Int Ed Engl 53, 1311–1315 (2014).

63. G. Shinar, E. Dekel, T. Tlusty, U. Alon, Rules for biological regulation based on error minimization. Proc Natl Acad Sci U S A 103, 3999–4004 (2006).

64. O. Brandman, T. Meyer, Feedback loops shape cellular signals in space and time. Science 322, 390–395 (2008).

65. X. Tan, N. Thapa, S. Choi, R. A. Anderson, Emerging roles of PtdIns(4,5)P2--beyond the plasma membrane. J Cell Sci 128, 4047–4056 (2015).

66. V. Marchesin, A. Castro-Castro, C. Lodillinsky, A. Castagnino, J. Cyrta, H. Bonsang-Kitzis, L. Fuhrmann, M. Irondelle, E. Infante, G. Montagnac, F. Reyal, A. Vincent-Salomon, P. Chavrier, ARF6-JIP3/4 regulate endosomal tubules for MT1-MMP exocytosis in cancer invasion. J Cell Biol 211, 339–358 (2015).

67. L. Mao, Y. Y. Zhan, B. Wu, Q. Yu, L. Xu, X. Hong, L. Zhong, P. Mi, L. Xiao, X. Wang, H. Cao, W. Zhang, B. Chen, J. Xiang, K. Mei, R. Radhakrishnan, W. Guo, T. Hu, ULK1 phosphorylates Exo70 to suppress breast cancer metastasis. Nat Commun 11, 117 (2020).

68. M. Uhm, M. Bazuine, P. Zhao, S. H. Chiang, T. Xiong, S. Karunanithi, L. Chang, A. R. Saltiel, Phosphorylation of the exocyst protein Exo84 by TBK1 promotes insulin-stimulated GLUT4 trafficking. Sci Signal 10, (2017).

69. D. Liu, X. Li, D. Shen, P. Novick, Two subunits of the exocyst, Sec3p and Exo70p, can function exclusively on the plasma membrane. Mol Biol Cell 29, 736–750 (2018).

70. E. J. Hartman, J. D. Romano, I. Coppens, The Rab11-Family Interacting Proteins reveal selective interaction of mammalian recycling endosomes with the <em>Toxoplasma</em> parasitophorous vacuole in a Rab11- and Arf6-dependent manner. bioRxiv, 2021.2006.2001.446625 (2021).

71. H. Kobayashi, M. Fukuda, Arf6, Rab11 and transferrin receptor define distinct populations of recycling endosomes. Commun Integr Biol 6, e25036 (2013).

72. T. Shiba, H. Koga, H. W. Shin, M. Kawasaki, R. Kato, K. Nakayama, S. Wakatsuki, Structural basis for Rab11-dependent membrane recruitment of a family of Rab11-interacting protein 3 (FIP3)/Arfophilin-1. Proc Natl Acad Sci U S A 103, 15416–15421 (2006).

73. T. D. Martin, X. W. Chen, R. E. Kaplan, A. R. Saltiel, C. L. Walker, D. J. Reiner, C. J. Der, Ral and Rheb GTPase activating proteins integrate mTOR and GTPase signaling in aging, autophagy, and tumor cell invasion. Mol Cell 53, 209–220 (2014).

74. G. Tang, L. Peng, P. R. Baldwin, D. S. Mann, W. Jiang, I. Rees, S. J. Ludtke, EMAN2: an extensible image processing suite for electron microscopy. J Struct Biol 157, 38–46 (2007).

75. D. N. Mastronarde, Automated electron microscope tomography using robust prediction of specimen movements. J Struct Biol 152, 36–51 (2005).

76. W. J. H. Hagen, W. Wan, J. A. G. Briggs, Implementation of a cryo-electron tomography tilt-scheme optimized for high resolution subtomogram averaging. J Struct Biol 197, 191–198 (2017).

77. J. R. Kremer, D. N. Mastronarde, J. R. McIntosh, Computer visualization of threedimensional image data using IMOD. J Struct Biol 116, 71–76 (1996).

78. A. Rizk, G. Paul, P. Incardona, M. Bugarski, M. Mansouri, A. Niemann, U. Ziegler, P. Berger, I. F. Sbalzarini, Segmentation and quantification of subcellular structures in fluorescence microscopy images using Squassh. Nat Protoc 9, 586–596 (2014).

79. S. J. Lord, K. B. Velle, R. D. Mullins, L. K. Fritz-Laylin, SuperPlots: Communicating reproducibility and variability in cell biology. J Cell Biol 219, (2020).

